# How the Heart Shapes the Mind: The Role of Cardiac Interoception in the Interaction between Autonomic Nervous Activity and Self-related Thoughts

**DOI:** 10.1101/2025.03.17.643435

**Authors:** Mai Sakuragi, Satoshi Umeda

## Abstract

Our thoughts often drift away from the tasks at hand. Various factors influence this phenomenon, including changes in the external environment, individual cognitive characteristics, and fluctuations in bodily responses. This study investigated the relationship between autonomic nervous fluctuations and thought state transitions, focusing on individual differences such as cardiac interoception. First, the heartbeat counting task was conducted, and the difference between the reported and actual number of heartbeats was used as an index of interoceptive accuracy. The participants then completed an auditory attention task while their cardiac activities were monitored. During the task, thought probes were randomly presented, and participants selected their thought content from eight categories and rated aspects such as task concentration and arousal. We estimated trial-by-trial thought states in a data-driven manner and examined how the current thought state, autonomic nervous activity, and individual cardiac interoceptive accuracy influenced the thought state in the next trial. The results demonstrated a strong association between higher cardiac interoceptive accuracy and the maintenance of similar states in subsequent trials when accelerated heart rates occurred during self-related thought states. Furthermore, the participants with higher depressive tendencies and interoceptive accuracy exhibited an increased likelihood of transitioning to self-related states when experiencing decreased heart rate during task-concentrated states. These results suggest that accurately detecting heart rate changes associated with specific thought states facilitates updates in first-person conscious experience, thereby biasing the transition patterns of subsequent thought states. This study provides new insights into the cognitive and physiological mechanisms underlying the dynamics of spontaneous thought.

## 1. Introduction

Mind-wandering (MW) is a phenomenon in which thoughts spontaneously drift away from the task at hand (Smallwood & Schooler, 2015). Early studies defined MW as task-unrelated thought and examined it by evaluating the task relevance of participants’ thoughts during laboratory tasks (Murray et al., 2020). In recent years, MW has been examined beyond task relevance, incorporating various components such as temporal orientation (Smallwood et al., 2009, 2011), arousal (Unsworth & Robison, 2018), emotional valence (Banks et al., 2016; Welhaf & Banks, 2024), self-relatedness (Babo-Rebelo et al., 2016, 2019), contemplation (Sakuragi et al., 2023, 2024), and intentionality (Seli et al., 2016). These elements are distinguishable based on participants’ self-reports and are associated with differences in task performance (Shrimpton et al., 2017), autonomic nervous activity (Ottaviani et al., 2013; Ottaviani, Medea, et al., 2015; Ottaviani et al., 2017), and neural activity (Babo-Rebelo, Richter, et al., 2016; Babo-Rebelo, Wolpert, et al., 2016; Bertossi & Ciaramelli, 2016). These findings suggest that thoughts previously categorized simply as task-related or task-unrelated may have encompassed diverse types of cognitive processes. Recent studies analyzing temporal dynamics of thought content have revealed the existence of complex states characterized by the coexistence of multiple thought components (Sakuragi et al., 2024; Shinagawa et al., 2023; Zanesco et al., 2020), indicating that various components constituting thoughts are not independent but interact dynamically to shape moment-to-moment thought states.

Extending this multidimensional understanding of MW, our previous studies have revealed that transition patterns of MW are associated with specific autonomic nervous activity patterns through interoception (Sakuragi et al., 2023; 2024). Interoception refers to the process by which the nervous system perceives, interprets, and integrates signals from within the body, particularly from visceral organs, operating at both conscious and unconscious levels (Khalsa et al., 2018). Our recent findings suggest that individuals with highly cardiac interoceptive accuracy are more accurate in detecting heart rate changes, which may lead to deep engagement in highly contemplative self-related thoughts (Sakuragi et al., 2023). Even during resting states, in individuals with accurate cardiac interoception, increased sympathetic activity is associated with the continuation of highly contemplative self-related thoughts. In contrast, increased parasympathetic activity is linked to transitions between lower-arousal thought states (Sakuragi et al., 2024). Previous research has demonstrated that interoceptive information facilitates emotional recognition and memory retrieval (Garfinkel et al., 2013; Garfinkel, Critchley, et al., 2016; Terasawa et al., 2013; Umeda et al., 2016). These cognitive experiences lead to updating first-person conscious experiences (Park & Tallon-Baudry, 2014; Tallon-Baudry et al., 2018) and may generate spontaneous transitions in thought. However, our research has only established the relationship between self-relatedness and contemplation of thought with heart rate and cardiac interoceptive accuracy, while the relationships with various other thought components mentioned in the previous chapter remain unclear. Furthermore, while we have primarily focused on examining the relationship between cardiac interoception, autonomic nervous activity estimated from heart rate, and thought processes, respiration is another critical bodily response related to interoception and thought. Previous research has demonstrated that cardiac and respiratory interoception can be dissociated (Garfinkel, Manassei, et al., 2016). In MW research, although MW has often been studied in the context of breath-focused meditation or breath-counting tasks (Hasenkamp et al., 2012; Lu & Rodriguez-Larios, 2022; Rodriguez-Larios & Alaerts, 2021), there has been little discussion regarding how respiration changes during the occurrence of MW and how individual differences in respiratory interoception might influence this process.

The dynamics of thought transitions during MW are influenced not only by autonomic nervous activity and interoception but also by various individual differences. These include clinical characteristics such as depression, anxiety, and attention deficit hyperactivity disorder (ADHD) symptoms (Bozhilova et al., 2020; Fell et al., 2023; Figueiredo et al., 2020; Ottaviani et al., 2016; Ottaviani, Shahabi, et al., 2015; Seli et al., 2019; Smallwood et al., 2007), cognitive traits such as worry and rumination (repetitively thinking about the causes, consequences, and symptoms of one’s negative affect; Nolen-Hoeksema, 1991), and behavioral tendencies such as attentional awareness in daily life (Belardi et al., 2022; Mrazek et al., 2012; Turkelson & Mano, 2022). Moreover, these individual differences are also reflected in specific patterns of autonomic nervous activity and interoceptive processing (Adams et al., 2022; Carney et al., 2005; Domschke et al., 2010; Gorman & Sloan, 2000; Hu et al., 2016; Kong et al., 2022; Paulus & Stein, 2010; Pollatos et al., 2009; Wang et al., 2013). Previous research has mainly focused on the relationship among depression or anxiety, autonomic nervous activity, and interoceptive processing. These two psychiatric disorders share common features of distorted interoception and excessive self-focused thinking (Paulus & Stein, 2010). However, it has been suggested that interoception may not be systematically impaired in these psychiatry tendencies (Jenkinson et al., 2024). Therefore, a comprehensive examination is needed regarding how autonomic nervous activity and interoception may influence the specific thought patterns associated with the aforementioned individual characteristics, including depression and anxiety.

Building on these findings, the present study aimed to elucidate the complex relationships among thought state transitions, autonomic nervous activity, and interoception through three primary objectives: (1) to estimate thought states by integrating multiple components of thought, (2) to examine the interaction among estimated thought state transitions, the fluctuations in autonomic nervous activity, and cardiac interoception, and (3) to investigate how individual differences in depression, anxiety, everyday attentional tendencies, and sensitivity to bodily sensations and emotions influence this relationship. The participants engaged in a simple auditory attention task while their electrocardiogram (ECG) and respiration were measured, and they regularly reported their thought contents during the task. We employed hidden Markov Models to estimate underlying thought states from time-series data of the participants’ thought components collected during the attention task. This model assumes the simultaneous presence of multiple components of thought (e.g., task concentration, temporal orientation, emotional valence) during MW, enabling the data-driven extraction of multiple thought states with distinct probability distributions for each component. Furthermore, this study focused on cardiac interoceptive accuracy (objective accuracy in detecting cardiac activity), a subcategory of interoception (Garfinkel & Critchley, 2013). To measure this index, the participants completed the heartbeat counting task (HCT; Schandry, 1981). Performance on this task represents how accurately individuals can perceive their heart rate. Individuals with higher performance on this task show more robust functional connectivity in interoception-related regions in fMRI studies (Chong et al., 2017) and greater amplitude of cardiac-related afferent signals in electroencephalography (Coll et al., 2021), even during the resting state. This suggests that the participants with higher cardiac interoceptive accuracy, as measured by the HCT, may have enhanced neural processing of interoception at an unconscious level, even when their attention is not directly focused on cardiac activity. Based on our previous findings (Sakuragi et al., 2023, 2024), we hypothesized that (1) sympathetic dominance would promote the persistence of self-related thoughts, while parasympathetic dominance would facilitate thought transitions, and (2) the influence of individual differences in thought-related characteristics on thought transition patterns would be mediated by autonomic activity and further modulated by interoceptive accuracy. For (2), we comprehensively and exploratorily examined which individual characteristics interact with autonomic nervous activity and interoception on thought transition patterns and how these interactions occur.

## 2. Material and Methods

### 2.1. Participants

Thirty-two healthy university students, graduate students, and working adults (11 males, 21 females; mean age 21 years, range 18-27 years) participated in the experiment. All participants had normal or corrected-to-normal vision. On the day of the experiment, the participants verbally confirmed that they had not taken any medications affecting mental or neurological functions, headache medication, cold medicine, or antihistamines; had no cardiac conditions; and had not consumed any vasoactive substances (including foods and beverages containing caffeine, alcohol, or nicotine) within three hours before the experiment. All participants provided written informed consent before participating in the experiment. After the experiment, the participants were debriefed and offered additional written consent for the use of collected data. This study was approved by the Research Ethics Committee of Keio University (approval number: 240040000) and conducted in accordance with the Declaration of Helsinki.

### 2.2. Apparatus

Throughout all experimental tasks, ECG was recorded using MP-150 and AcqKnowledge (Biopac Systems, Santa Barbara, CA), while continuous blood pressure was measured using VitalStream (Caretaker Medical, Charlottesville, VA). ECG measurements were obtained with electrodes attached to the right hand and both feet, and continuous blood pressure was monitored via a cuff attached to the middle finger of the right hand. During the sound attention task, in addition to ECG and continuous blood pressure measurements, thoracic and abdominal circumference changes due to respiration were recorded using a respiratory measurement belt (RESP100C, Biopac Systems, Santa Barbara, CA).

### 2.3. Questionnaire

The questionnaires used in this study were the State-Trait Anxiety Inventory (STAI; Spielberger et al., 1970; Spielberger & Gorsuch, 1983), Beck Depression Inventory-II (BDI-II; Beck, 1961), Body Perception Questionnaire (BPQ-BA; Cabrera et al., 2018, Kobayashi et al., 2021), Multidimensional Assessment of Interoceptive Awareness Japanese version (MAIA-J; Shoji et al., 2018), Mind-Wandering Questionnaire (MWQ; Kajimura & Nomura, 2016), Daydream Frequency Scale (DDFS; Kajimura & Nomura, 2016), Mindful Attention Awareness Scale (MAAS; Brown & Ryan, 2003; Fujino et al., 2015), Five Facet Mindfulness Questionnaire (FFMQ; Baer et al., 2006; Sugiura et al., 2012), and Rumination-Reflection Questionnaire (RRQ; Takano & Tanno, 2008, Trapnell & Campbell, 1999). The STAI-State was administered before and after the auditory attention task, while the participants completed all other questionnaires at home after the experimental tasks were finished.

### 2.4. Procedure

#### 2.4.1. Heartbeat Counting Task (HCT)

While monitoring ECG and continuous blood pressure, we conducted a heartbeat counting task (HCT) to assess interoceptive accuracy (Schandry, 1981). The task consisted of three components: resting pulse measurement, the HCT, and the time estimation task (TET). During these tasks, the participants placed their left arm on the table to report heartbeat counts while maintaining their right hand (equipped with the continuous blood pressure monitor) at rest on their right knee. For the heartbeat counting task, the participants silently counted their heartbeats during specified intervals (25 seconds × 2, 35 seconds × 2, 45 seconds × 2) and reported their count after each interval. The participants were instructed to avoid counting heartbeats through physical manipulation (touching their body, pressing against furniture) or breath holding. Following modified instructions (Desmedt et al., 2018), the participants were directed not to estimate imperceptible heartbeats. After each trial, the participants rated their confidence on a scale of 1 to 10, with higher scores indicating greater perceived accuracy between reported and actual heartbeat counts. In the TET, the participants subjectively estimated and reported the duration of fixed time intervals (23 seconds × 2, 49 seconds × 2, 56 seconds × 2). The intervals for both tasks were administered in randomized order.

#### 2.4.2. Sound Attention Task

After completing the HCT, the participants completed the STAI-State questionnaire to record their mood. Following this, respiratory belts were attached to their chest and abdomen, as well as the ECG and continuous blood pressure monitors. The participants were then seated comfortably in a soundproof laboratory. We instructed them to keep their eyes open, maintain their right hand at rest on their knee, and keep both feet flat on the floor to ensure high signal quality. Five minutes of baseline physiological data were collected before task initiation.

During the task, the participants maintained their gaze on a central fixation cross while listening to auditory stimuli. The sound stimuli were 1000 Hz, 65 dB pure tones, always presented at a tempo equal to the median RR interval of each participant’s resting heart rate before the task. The interval between sound stimuli was kept constant to prevent the acceleration or deceleration of sound stimuli from influencing the heart rate tempo during the task. The participants responded as quickly and accurately as possible by pressing the nine key with their right hand when they heard the sound. Following thought probes were presented every 50-70 key presses (randomly determined between 50-70 from a uniform distribution) to assess the participants’ thought states. The flow of the task per trial is summarized in Fig. 1.

**Fig. 1.**
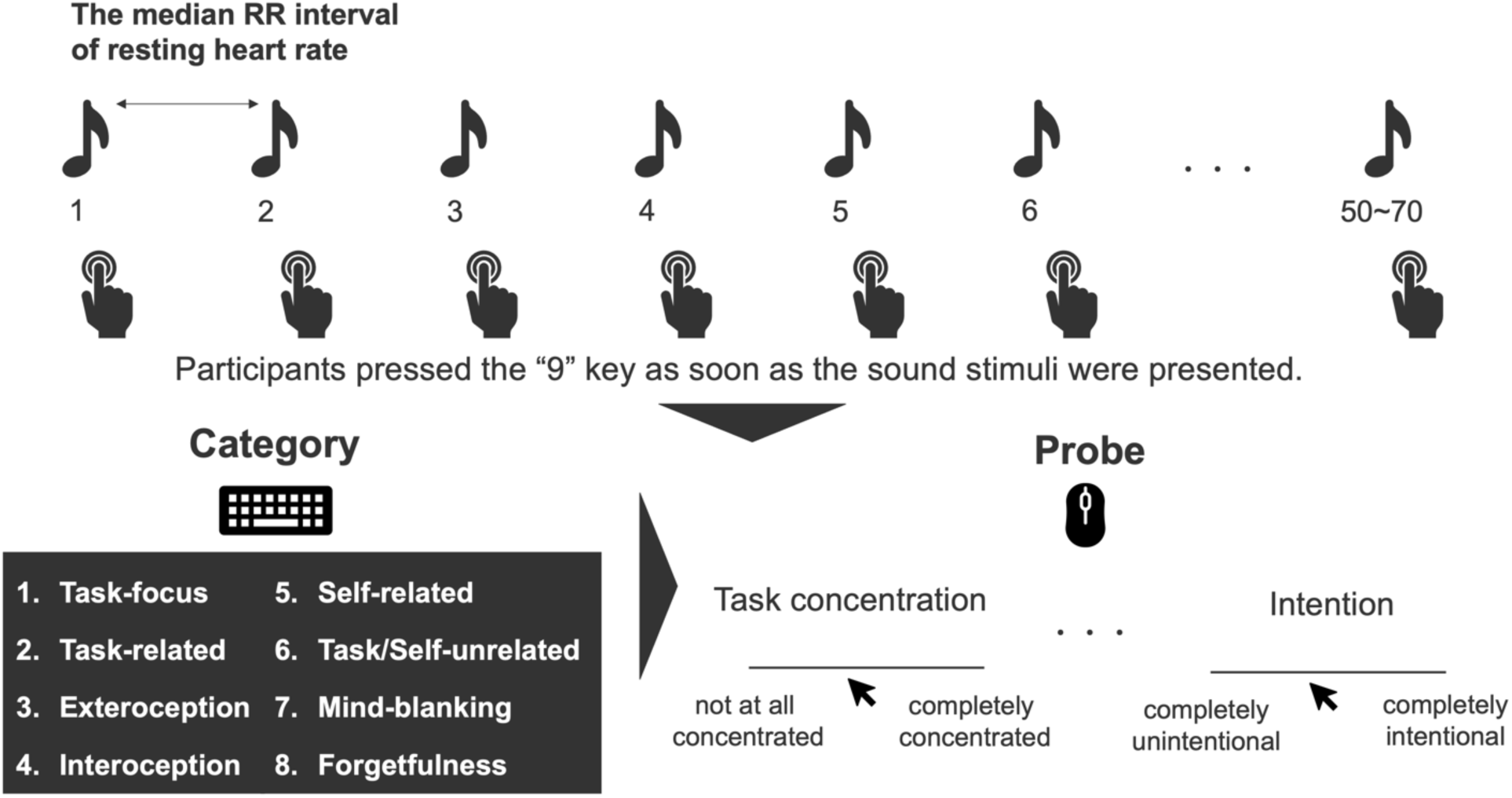
Experimental Task Procedure. The participants performed a task requiring them to press a keyboard key each time they heard an auditory stimulus presented at a constant rate. The auditory stimuli were presented at intervals matching the median heart rate measured during the preceding resting state. After 50-70 key presses, thought probes were presented, asking about the participant’s thought state immediately preceding the probe. The participants first selected one of eight categories that best represented their thoughts or occupied most of the task time by pressing the corresponding key. Based on their category selection, additional probes were presented asking about various components of their thought (see Fig. 2), to which the participants responded by clicking the appropriate position on a visual analog scale displayed on the screen.

When presented with thought probes, the participants first selected their predominant thought content, defined as either the content that occupied the most extended duration or left the strongest impression, from eight categories: 1. task-focus, 2. task-related (concerning elapsed time or one’s performance in the experiment), 3. exteroception (environmental changes unrelated to the task), 4. interoception (bodily sensations), 5. self-related (personal past episodes or future plans), 6. task/self-unrelated thoughts (matters neither directly related to the task nor the individual, such as general knowledge), 7. mind-blanking (absence of thought content), and 8. forgetfulness. The participants responded by pressing the corresponding numerical key.

After the participants selected categories 1-6 in the initial thought category question, they evaluated various components of their thoughts using visual analog scales (VAS; 0-100). A line segment was presented on the screen with labels at both ends and, when necessary, in the middle section (as described below). The participants responded by using their left hand to operate the mouse and clicking at the appropriate position on the line. The participants evaluated the following nine components of their thoughts: task concentration (Probe1; 0: “not at all concentrated” to 100: “completely concentrated”), time (Probe2; 0: “past”, 50: “present”, 100: “future”), arousal (Probe3; 0: “very calm” to 100: “very excited”), valence (Probe4; 0: “negative”, 50: “neutral”, 100: “positive”), agency (Probe5; 0: “no sense of agency” to 100: “very strong sense of agency”), self-focus (Probe6; 0: “no self-focus” to 100: “intense self-focus”), bodily information (Probe7; 0: “no bodily information” to 100: “abundant bodily information”), contemplation (Probe8; 0: “no contemplation” to 100: “very deep contemplation”), and intention (Probe9; 0: “completely unintentional” to 100: “completely intentional”). For the time probe, 0 represented the most distant imaginable past, 50 represented the present moment, and 100 represented the most distant imaginable future. Time, valence, agency, and self-focus were adopted based on previous research examining the possibility that self-relevant content during spontaneous thought is encoded in conjunction with the neural processing of cardiac signals in the default mode network (Babo-Rebelo, Richter, et al., 2016). The agency and self-focus measures were originally referred to as the “I” and “Me” scales; however, we modified these expressions when creating Japanese instructions. This paper describes the English translations of our modified Japanese terms. The agency assessed the participant’s engagement as the protagonist or the agent in the thought, while self-focus evaluated the extent to which thought content was oriented toward oneself. Based on previous research indicating that intentional and unintentional mind wandering are dissociable cognitive experiences with distinct mechanisms, we employed an intention scale to assess the degree to which the participants intentionally engaged in their reported thoughts (Seli et al., 2016). The participants did not respond to all probes each time; probes that were self-evident from the definition of the initially selected category were omitted. Specifically, time, valence, agency, self-focus, and bodily information probes were sometimes omitted depending on the category initially chosen by the participant, and specific values were interpolated during subsequent data processing. For example, if a participant selected the task-focus category, it was self-evident that they were thinking about the current state during the task, so the time probe was interpolated with a value of 50, corresponding to the present, during post-experimental data processing. When the participants selected “mind-blanking” or “forgetfulness,” responses to questions about thought components were coded as a separate “no response” category. Fig. 2 summarizes which probes were presented following responses to each category and which values were used for interpolation when probes were omitted.

**Fig. 2.**
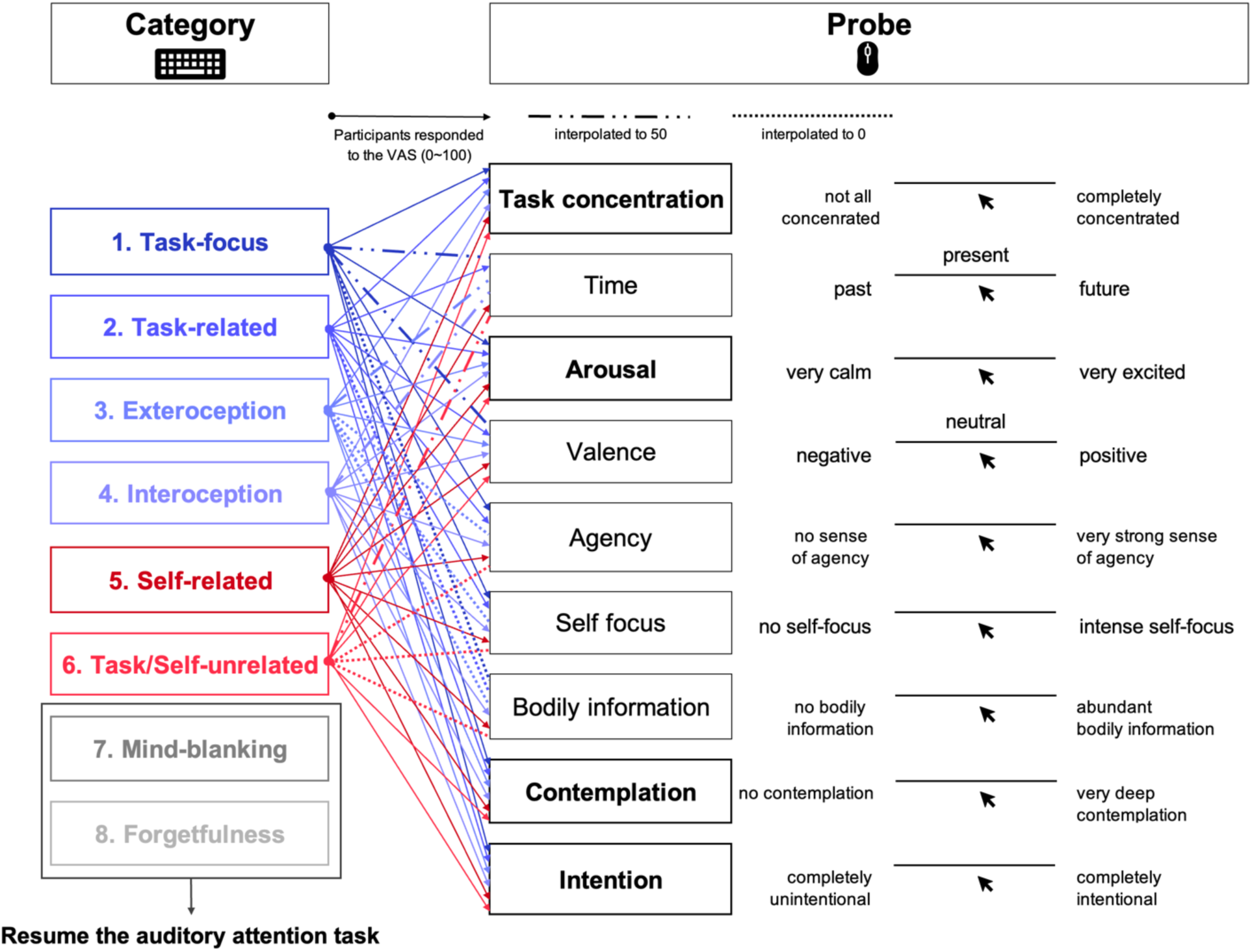
Thought Categories Selected by the Participants and Corresponding Probes for Each Category The participants first selected one of eight categories that best represented their thoughts or occupied the majority of task time by pressing the corresponding keyboard key. Next, questions about various components of thought (connected to each category labeled by arrows) were presented based on their selected category. The participants responded to probes by clicking the appropriate position on the solid line on the screen using a mouse with their left hand. Scores were quantified as zero at the left end of the line, 50 at the center, and 100 at the right end. If the participants selected “mind-blanking” or “forgetfulness,” the thought probe ended, and the auditory attention task resumed. Probes in bold text and surrounded by thick borders were always presented when any category other than “mind-blanking” and “forgetfulness” was selected. Questions about components that were self-evident from the definition of each category were not presented during the task, and appropriate values were interpolated during subsequent data processing. Probes connected to category labels by dashed lines were interpolated with a value of 50, while those connected by dotted lines were interpolated with a value of 0. When the participants selected “mind-blanking” or “forgetfulness,” responses to questions about thought components were coded as a separate “no response” category and used in subsequent analyses.

The duration per trial varied depending on the participants’ resting heart rate intervals and response speed to thought probes, but typically ranged from approximately 1.5 to 2 minutes. The task comprised 15 trials in total, with the entire task lasting 25-30 minutes. After the task, the participants completed the STAI-State questionnaire assessing post-task mood state. Additionally, we conducted an original questionnaire to investigate thought patterns in daily life during the week before the experiment. As we do not report the response data from this original questionnaire in the present paper, the questionnaire’s content is summarized in the Appendix.

### 2.5. Pre-Processing

#### 2.5.1. The score of the HCT

Based on the actual heartbeat count recorded by AcqKnowledge and self-reported heartbeat counts, performance scores for each trial were calculated using the formula: 1 - |actual heartbeats – reported heartbeats|/actual heartbeats. For each participant, the mean score across six trials served as an index of cardiac interoceptive accuracy. Similarly, confidence ratings were averaged across the six trials to measure each participant’s confidence in their heartbeat perception ability. For the TET, the score was calculated using the formula 1 - |actual time-lapse - reported time lapse|/actual time-lapse. To confirm that the HCT scores were not affected by inferences based on time estimates (Desmedt et al., 2020), we calculated Pearson’s product-moment correlation coefficient for the HCT and the TET scores.

#### 2.5.2. Electrocardiogram and Respiration

R-waves in the ECG data were detected using AcqKnowledge. Raw data preprocessing and respiratory phase detection were performed through the BreathMetrics toolbox. Before decomposing raw respiratory flow recordings into phase data, measurement noise, and signal drift were removed. Initially, signals underwent mean smoothing with a 25-millisecond window. Overall linear drift was eliminated by subtracting the slope of the data’s linear regression model. Local signal drift was corrected against a continuous one-minute sliding average baseline window, followed by padding removal. Subsequently, the respiratory phase at each time point was detected based on the processed respiratory data. This study collected respiratory data from both thoracic and abdominal sites. For each participant, the respiratory data that yielded more detected cycles between thoracic and abdominal measurements was selected for subsequent analyses.

#### 2.5.3. Thought Probes

The distribution of VAS values for each thought probe is summarized in Fig. 3. As a result of value substitution based on the participants’ category responses (see 2.4.2.), responses were concentrated at the midpoint of 50 for time and valence, and at 0 for agency, self-focus, and bodily information. These five probe datasets were not used in subsequent thought state extraction to prevent any potential bias from this post-hoc processing-induced distortion in response distributions (see 2.6.2.). Considering the data distribution of each probe, responses were categorized into four levels: for time and valence, values below 30 were coded as 1, 30- 50 as 2, 51-70 as 3, and above 70 as 4; for agency and self-focus, values below 25 were coded as 1, 25-50 as 2, 51-75 as 3, and above 75 as 4. Regarding bodily information, responses were highly skewed toward either 0 or 100, with values approaching 100 when the initial thought category response was 4 (interoception). Therefore, we determined that the VAS responses for this probe contained minimal additional information and excluded it from all subsequent analyses. In the following model analysis, we utilized data from task concentration, arousal, contemplation, and intention, in addition to category (probes highlighted in orange frames in Fig. 3).

**Fig. 3.**
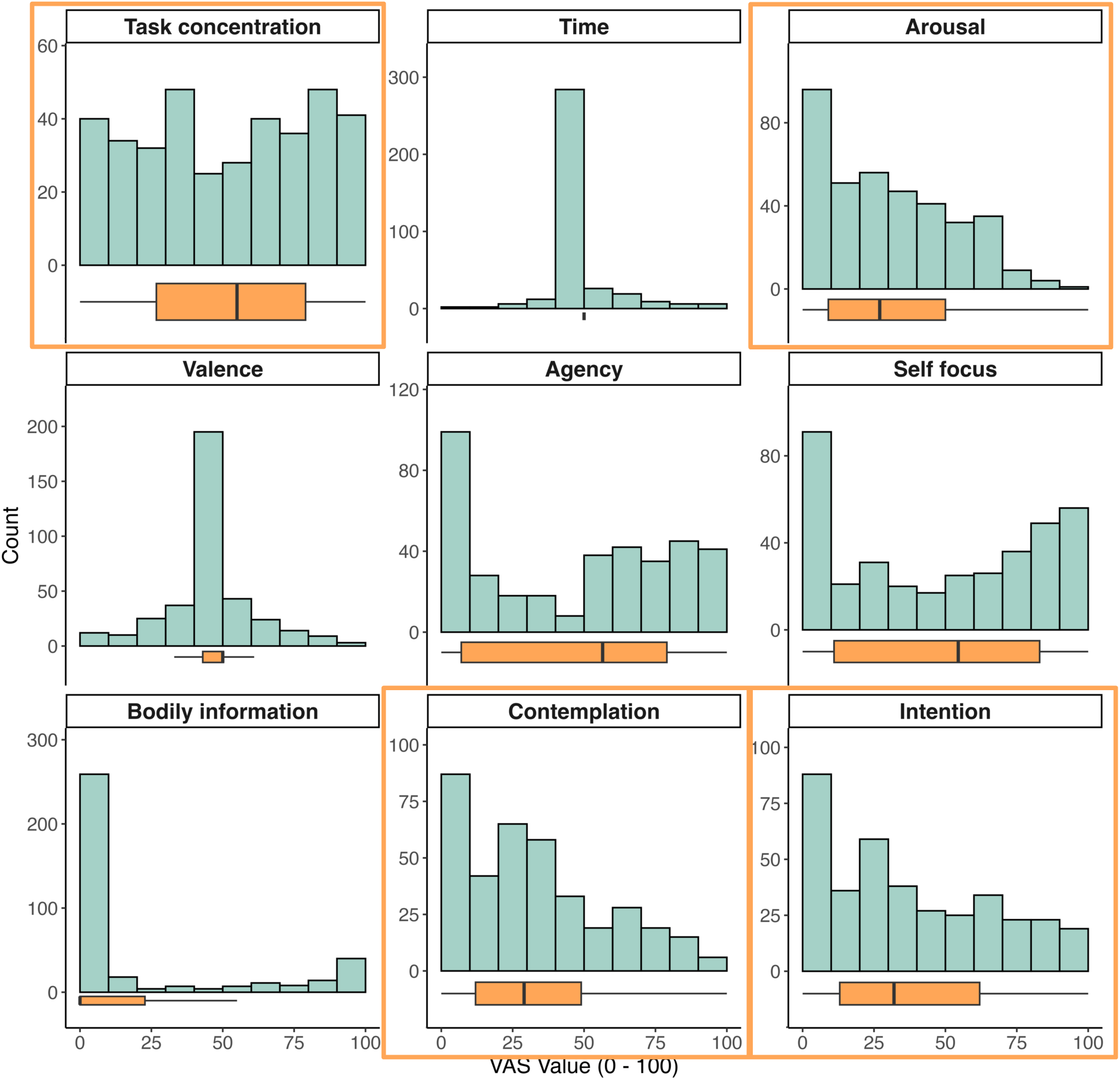
Distribution of Visual Analogue Scale (VAS) Scores for Each Thought Probe. This figure summarizes VAS responses to thought probes presented during the Sound Attention Task (Section 2.4.2). The x-axis represents VAS values (0-100), while the y-axis shows the frequency of responses. The box plots below the histogram show the minimum and maximum VAS values at both ends, with the first and third quartiles at the box edges, and the median value in the center of the box. Concentration (Probe1; 0: “not at all concentrated” to 100: “completely concentrated”); Time (Probe2; 0: “past”, 50: “present”, 100: “future”); Arousal (Probe3; 0: “very calm” to 100: “very excited”); Valence (Probe4; 0: “negative”, 50: “neutral”, 100: “positive”); Agency (Probe5; 0: “no sense of agency” to 100: “very strong sense of agency”); Self-focus (Probe6; 0: “no self-focus” to 100: “intense self-focus”); Bodily information (Probe7; 0: “no bodily information” to 100: “abundant bodily information”); Contemplation (Probe8; 0: “no contemplation” to 100: “very deep contemplation”); Intention (Probe9; 0: “completely unintentional” to 100: “completely intentional”). The thought probes highlighted in orange frames were used in the subsequent analysis (Section 2.6.2).

#### 2.5.4. The Score of the Questionnaires

For all questionnaires collected in this study, total scores for STAI, BDI, BPQ, MWQ, DDFS, and MAAS were calculated by averaging all items after accounting for reverse-scored items. For MAIA, mean scores were calculated for the following subscales: Noticing ((Q1+Q2+Q3+Q4)/4), Not-Distracting ((Q5+Q6+Q7)/3), Not-Worrying ((Q8+Q9+Q10)/3), Attention Regulation ((Q11+Q12+Q13+Q14+Q15+Q16+Q17)/7), Emotional Awareness ((Q18+Q19+Q20+Q21+Q22)/5), Self-Regulation ((Q23+Q24+Q25+Q26)/4), Body Listening ((Q27+Q28+Q29)/3), and Trusting ((Q30+Q31+Q32)/3). For RRQ, mean scores were calculated separately for its two subscales, Rumination and Reflection, each consisting of 12 items.

### 2.6. Analysis

#### 2.6.1. Indicators of Autonomic Nervous Activity

We analyzed R-wave intervals (RR intervals) from ECG data to measure changes in heart rate. For each participant, RR interval data were standardized after excluding periods during probe responses, and mean standardized RR intervals were calculated for each trial. Additionally, we computed the Root Mean Square of Successive Differences (RMSSD) for each trial using non-standardized RR interval data by calculating the square root of the mean squared differences between successive heartbeats. RMSSD enables quantification of autonomic nervous activity over relatively short periods, such as 10 seconds (Nussinovitch et al., 2011).

Regarding the respiratory data, one respiratory cycle was defined as a combination of one expiratory and one inspiratory phase. The number of respiratory cycles per trial was calculated for each participant and standardized to cycles per 60 seconds for the analysis (respiration rate).

#### 2.6.2. Hidden Markov Model

Hidden Markov Models (HMM) were employed to estimate thought states generated from temporal transition patterns of thought content and components. This model assumes observable data is generated by a sequence of unobservable states, where each hidden state has specific transition probabilities between states and probabilities of generating observable outputs. This approach has been previously utilized to reveal behavioral and attentional states underlying thought experience sampling data and task response times (Bastian & Sackur, 2013; Sakuragi et al., 2024; Shinagawa et al., 2023; Zanesco et al., 2020).

We used time series data (15 probes each) of thought content and four thought probes, task concentration (Probe1), arousal (Probe3), contemplation (Probe8), and intention (Probe9) as HMM inputs. These four thought probes were presented regardless of which category the participants reported, so we determined they were optimal for extracting thought states based on the overall response patterns across the participants. These four probes were divided into four levels based on their quartile points (Probe1; level1: below 26.8, level2: 26.8-55, level3: 55-79, level4: above 79, Probe3; level1: below 9, level2: 9-27, level3: 27-50, level4: above 50, Probe8; level1: below 12, level2: 12-29, level3: 29-49, level4: above 49, Probe9; level1: below 13, level2: 13- 32, level3: 32-62, level4: above 62). Trials with thought content categorized as 7 or 8 were assigned to the fifth level because data for these four probes were missing.

HMM was implemented using the depmixS4 package in R 4.3.2, which employs expectation-maximization algorithms to perform maximum likelihood estimation on datasets with hidden variables, estimating HMM parameters, including state transitions, outputs, and initial state probabilities. Following previous research, the number of hidden states was determined using Bayesian Information Criterion (BIC), a model selection criterion balancing model complexity and data fit (Shinagawa et al., 2023; Zanesco et al., 2020). To enhance model stability and reproducibility, the estimation of the selected model was repeated 100 times, and the model with the lowest BIC was adopted. Based on the adopted model, we presented the response distribution of thought content and contemplation in each thought state, along with state occurrence probabilities and transition probabilities between states.

#### 2.6.3. Characteristics of each Thought States

We compared the following variables across thought states estimated by HMM: reporting probabilities, VAS values of thought probes used in HMM estimation (Probes 1, 3, 8, and 9), reporting probabilities for each level of thought probes not used in HMM estimation (Probes 2, 4, 5, and 6), RR intervals, RMSSD, and respiratory rates. For the statistical analysis, we constructed linear mixed models with different specifications: For reporting probabilities, RR intervals, RMSSD, and respiratory rates, thought state (States 1, 2, and 3) was set as the independent variable. For VAS values of Probes 1, 3, 8, and 9, the thought state (States 1 and 2) was the independent variable. For reporting probabilities of Probes 2, 4, 5, and 6, both main effects and interactions between thought state (States 1 and 2) and level (levels 1-4) were included as independent variables. Random effects structures varied across models: participant-only random effects were included for reporting probabilities and Probes 2, 4, 5, and 6 for reporting probabilities, while both participant and trial numbers were included as random effects for VAS values of Probes 1, 3, 8, and 9, RR intervals, RMSSD, and respiratory rates.

Bayesian approach was used to estimate the parameters of these statistical models. We used the brms R package to analyze (Bürkner, 2017, Bürkner, 2018) and estimated the parameters using four Markov chain Monte Carlo chains. For each of the four chains, we performed 2000 iterations, with the first 1000 iterations used as burn-in samples. This resulted in 1000 posterior samples per chain, which were combined across the four chains to obtain 4000 posterior samples for each parameter. For prior distributions, we used non-informative flat priors for fixed effects parameters and Student’s t-distributions centered at 0 with 3 degrees of freedom and a scale of 2.5 for intercepts. For random effects, we specified LKJ prior distributions (parameter = 1) for correlation matrices and Student’s t-distributions centered at 0 with 3 degrees of freedom and a scale of 2.5 for standard deviation components. Model convergence was evaluated based on the Gelman-Rubin convergence statistic R^ (Gelman & Rubin, 1992), with values close to 1 indicating negligible differences in within-chain and between-chain variance. The mean and 95 % confidence interval (CI) of the parameter estimates were used to examine the effect of each parameter. In this study, we employed the region of practical equivalence (ROPE) to examine the extent to which each parameter has an effect based on the estimated coefficients. The ROPE is a range within which a parameter can be considered to have no practical effect. We assessed the degree to which each parameter affected the dependent variable by determining the extent to which the credible interval of the estimated value for each parameter was within the ROPE. In this study, following Kruschke & Liddell (2018), we defined the ROPE as the range from −0.1* the dependent variable’s SD of posterior predicted value to 0.1* the dependent variable’s SD of posterior predicted value. When the credible interval of an estimated variable lies entirely within the ROPE, the parameter is deemed to have no practical effect. Conversely, if the credible interval fully surpasses the ROPE, the parameter is considered to have a practical effect. In cases where the credible interval partially overlaps the ROPE, the practical effect of the parameter remains uncertain.

To examine the relationship between thought states during the experiment and individual characteristics, we calculated Pearson’s correlation coefficients between the probability of occurrence for each thought state and questionnaire scores measuring daily thought tendencies (MWQ, DDFS, MAAS, STAI, BDI, RRQ_rumination, RRQ_reflection, and BPQ) and interoceptive ability (The HCT scores and BPQ).

#### 2.6.4. Multinomial Logistic Regression Model

In Section 2.6.3, we examined the relationship between autonomic nervous activity associated with thought states at each time point and the relationship between individual characteristics of participants and the reporting probability of each thought state. Additionally, considering that both thought and autonomic nervous activity exhibit dynamic changes over time, we sought to investigate how fluctuations in autonomic nervous activity influence changes in thought states. Therefore, we estimated and compared three multinomial logistic regression models to investigate the relationships between transition probabilities among three thought states between consecutive trials, autonomic nervous activity, and individual differences in interoceptive accuracy. We included interoceptive accuracy in this analysis because our previous research has revealed that the relationship between thought state transition patterns and autonomic nervous activity differs depending on interoceptive accuracy. In the current study’s model, we used the thought state and indicators of autonomic nervous activity (RR, RMSSD, or respiratory rate) at trial t, along with individual participants’ cardiac interoceptive accuracy (HCT scores), as independent variables, and the thought state at trial t+1 as the dependent variable. Based on the hypothesis that differences in the ability to accurately perceive changes in autonomic nervous activity associated with specific thought states would lead to different patterns of subsequent thought transitions, we focused on the interaction between thought state, indicators of autonomic nervous activity, and HCT scores. For each participant, we considered the individual differences in how the thought state × RR × interoceptive accuracy interaction affects the thought state of the subsequent trial (random slopes) while controlling for trial-by-trial variation as random intercepts. The dependent variable was assumed to follow a categorical distribution with a logit link function. The widely applicable information criterion (WAIC; Watanabe, 2010) was used to determine which model to adopt. This indicator estimates the expected log pointwise predictive density for a new dataset that integrates the posterior distributions of the model parameters, allowing for a combined assessment of model fit and complexity. Because a lower WAIC value indicates that the model fits the observed data better, a model with a lower WAIC value was adopted. The Bayesian model specification was identical to that described in Section 2.6.3.

#### 2.6.5. Relationships Between Individual Components, Autonomic Activity, and Thought Pattern Probabilities During Experiments

Furthermore, we aimed to elucidate how the interactive relationship between thought transition patterns and autonomic nervous activity during the experiment varies not only with interoceptive accuracy but also with individual differences in attentional components, mindfulness in daily life, and tendencies toward psychiatry. For each participant, we utilized data including the probability of each thought transition pattern (e.g., State1→1, State1→2…) calculated from the posterior predictive probabilities estimated from the model in 2.6.4., the mean RR interval corresponding to each thought transition pattern, interoceptive accuracy, and questionnaire scores measuring individual differences in thought-related components (MWQ, DDFS, MAAS, STAI, BDI, RRQ_rumination, and RRQ_reflection).

First, we calculated Pearson’s correlation coefficients between the probability of each thought transition pattern and questionnaire scores. For thought transition patterns that showed significant correlations with questionnaire scores, we conducted structural equation modeling (SEM) to examine the effect of individual traits, as indicated by questionnaire scores, on the probability of thought transitions. In this model, questionnaire scores influenced both RR intervals, an indicator of autonomic nervous activity that showed the influence on thought transition patterns in the multinomial logistic regression model, and interoceptive accuracy, which in turn mediated their effects on thought transitions. Additionally, interoceptive accuracy was modeled as a moderator, influencing the relationship between RR intervals and the probability of thought transitions.

The SEM model was specified as follows:

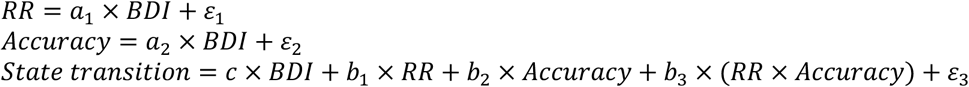

The indirect effects were calculated as:

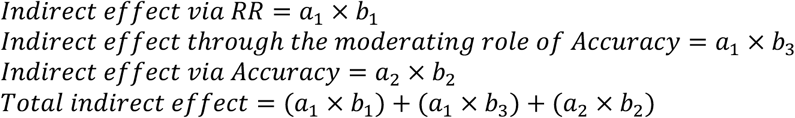

The total effect was defined as follows:

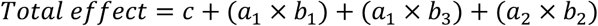

All variables included in the model were standardized, and 1,000 bootstrap resamples were performed. This analysis allowed us to examine not only the direct and indirect effects of individual traits on thought transitions but also how interoceptive accuracy modulates the influence of RR intervals on these transitions.

## 3. Results

### 3.1. Estimated Thought States and Their Characteristics

HMM was employed to estimate the thought states underlying self-reports of thought content, task concentration, arousal, contemplation, and intention. A three-state model was adopted as it yielded the smallest BIC value (1: 7898.829, 2: 6175.614, 3: 5922.905, 4: 6026.713, 5: 5980.557, 6: 6196.458, 7: 6282.750, 8: 6575.694). The distribution of thought content across estimated states (Categories 1-8; see Section 2.4.2) is shown in Fig. 4A. In State 1, task focus showed a higher selection probability than other thought contents. Conversely, in State 2, the probability of selecting task focus decreased, with self-related thoughts being most frequently selected. In State 3, the participants reported either a complete absence of thoughts or an inability to recall their previous thought content. Fig. 4B illustrates the distribution of VAS values for thought probes (Probes 1, 3, 8, 9) used in the HMM estimation across estimated thought states. Linear mixed model estimation revealed significant differences in VAS values between States 1 and 2 for task concentration, arousal, and contemplation (Probe 1, 3, 8) (Task concentration: *b* = −28.314, 95% CI [-33.720; −22.994]; Arousal: *b* = 21.866, 95% CI [17.220; 26.442]; Contemplation: *b* = 18.174, 95% CI [12.785; 23.635]). Task concentration was higher in State 1, while arousal and contemplation were higher in State 2. These results suggest that in State 1, although the participants’ thoughts were dominated by task focus, both arousal and contemplation levels were low, indicating a somewhat passive engagement with the task. In contrast, in State 2, the participants rarely focused on the task but were deeply engaged in self-related thoughts. Based on the distribution of selected thought content and response for thought probes, these three thought states were defined as on-task, highly contemplative self-related, and thought absence, respectively. Furthermore, Fig. 4D shows the reporting probabilities for each level of thought probes not used in the estimation (Probes 2, 4, 5, 6). Linear mixed model estimation revealed significant interactions between thought states (States 1, 2) and level only for valence (valence: *b* = −0.409, 95% CI [-0.680; −0.143]). The distribution of VAS values revealed that State 1 had a higher probability of level 2 responses (VAS values 30-50) than State 2. Level 2 represents near-neutral emotional valence. Although not statistically significant, State 2 tended toward higher proportions of level 1 and 4 responses, indicating strongly negative or positive emotional valence. These results demonstrated that self-relevant thoughts (State2) were associated with more pronounced emotional valence than State1.

**Fig. 4.**
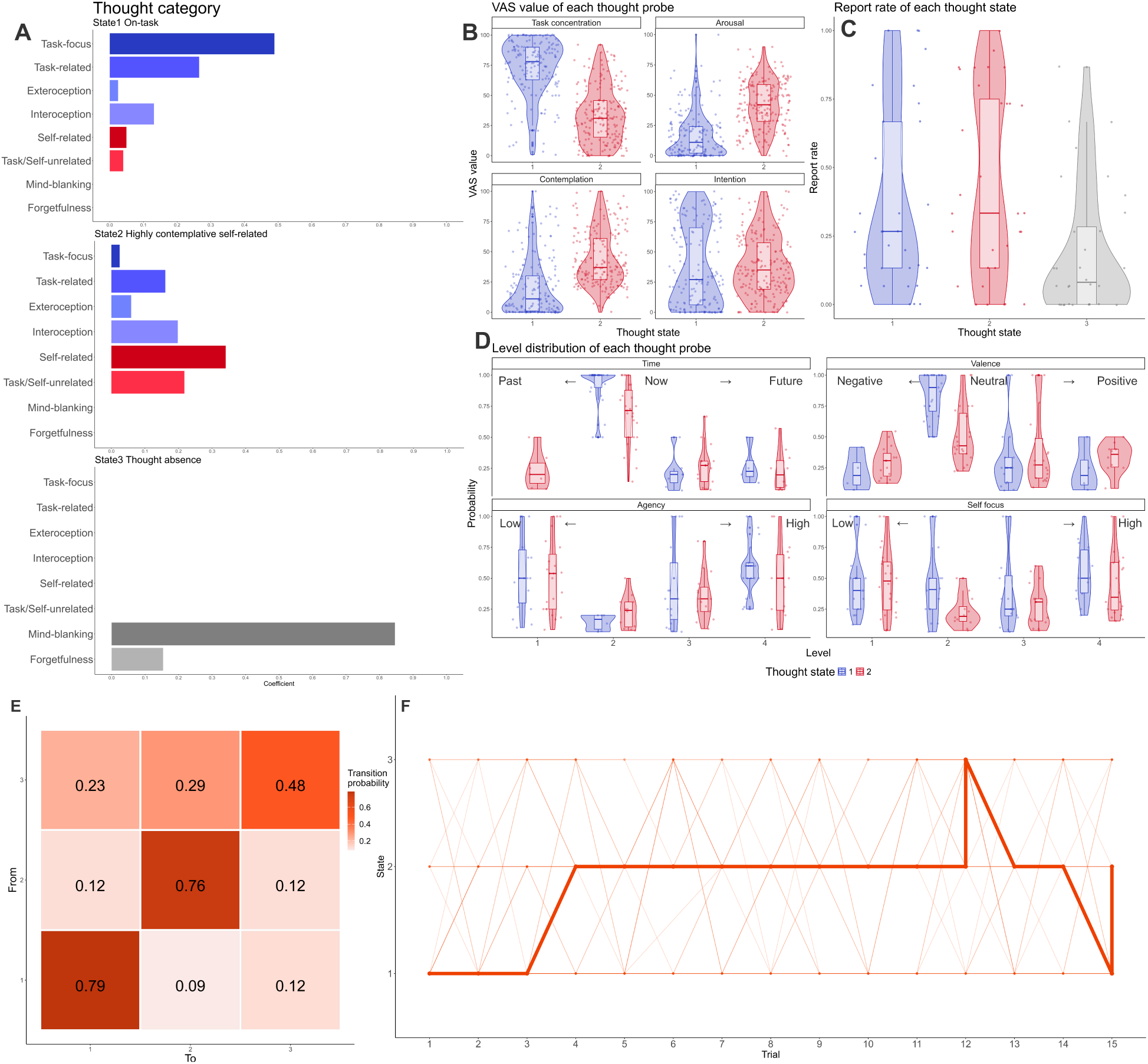
Information about Thought States Estimated by HMM (A) Distribution of thought categories within each thought state. The y-axis shows the names of thought categories, and the x-axis indicates the probability of each thought category being reported in respective thought states. (B) Distribution of VAS values for task concentration (Probe 1), arousal (Probe 3), contemplation (Probe 8), and intention (Probe 9) across thought states. The x-axis represents thought state numbers, and the y-axis shows VAS values (0-100). VAS values for task concentration, arousal, and contemplation showed statistically significant differences between States 1 and 2. Based on the distributions of thought categories and other components shown in (A) and (B), States 1-3 were labeled as on-task, highly contemplative self-related, and thought absence, respectively. (C) Distribution of report rates for each thought state. The x-axis shows thought state numbers, and the y-axis indicates the probability of each participant reporting respective thought states. Significant differences in report rates were observed only between States 1 and 3. (D) Distribution of reporting probabilities across levels for time (Probe 2), valence (Probe 4), agency (Probe 5), and self focus (Probe 6) in each thought state. Levels represent quartile divisions of VAS value distributions for each Probe. The x-axis shows level values, the y-axis indicates reporting probabilities, and colors denote different States. Only the reporting probability for level 2 of Valence (emotionally neutral thoughts) showed significant differences across States. In (B), (C), and (D), distributions along the y-axis are represented by violin and box plots, with individual data points showing participant-specific values. (E) Transition and continuation probabilities between thought states. The y-axis indicates the initial thought state, and the x-axis shows the destination thought state. For all thought states, continuation within the same state was more likely than transitions between different states. Among transitions between different states, the most frequent was from State 2 (highly contemplative self-related) to State 3 (thought absence). (F) The most frequently reported thought states were in each trial. The x-axis shows trial numbers, and the y-axis indicates thought state numbers. The thickest line connects the most frequently reported states across trials, while the lighter background lines represent the participants’ thought states for each trial. As a general trend, the participants were initially on-task (State 1) at the beginning of the task, became more likely to report self- related (State 2) during the middle phase, and showed instances of thought absence (State 3) toward the end of the task.

Comparison of report rates across the three thought states using linear mixed models revealed significant differences only between on-task (State 1) and thought absence (State 3) (*b* = −0.177, 95% CI [-0.336; −0.024]), suggesting that thought absence state occurred less frequently than task-focused state (Fig. 4C). Fig. 4E depicts the transition probabilities between estimated thought states, and Fig. 4F shows temporal changes in the most frequently reported thought states. All thought states showed the highest transition probabilities between consecutive trials of the same state, indicating state persistence across successive trials. Among transitions between different states, the transition from thought absence (State3) to highly contemplative self-related (State2) exhibited the highest probability. Temporal changes in predominant thought states across trials revealed that on-task (State 1) were most frequent during initial trials, while highly contemplative self-related (State 2) became predominant after the fourth trial. Thought absence state (State 3) tended to occur more frequently in the latter half of the task. These findings suggest that the participants initially maintained task concentration but gradually became immersed in vivid self-related thoughts and, in some cases, experienced thought absence states as their consciousness level decreased.

We examined whether individual characteristics, including interoceptive accuracy, daily thought tendencies, and mental health indicators, influenced the frequency of reporting specific thought states. We calculated correlation coefficients between scores of each index (Accuracy, STAI, BDI, BPQ, MAIA-J, MWQ, DDFS, MAAS, FFMQ, RRQ) and the individual reporting probabilities for each thought state. The analysis revealed marginally significant correlations between on-task (State 1) reporting probability and MAAS, MWQ, and RRQ_rumination scores (MAAS: *r* = −0.422, *p* = 0.077; MWQ: *r* = −0.407, *p* = 0.077; RRQ_rumination: *r* = −0.412, *p* = 0.077) while showing marginally significant positive correlations with highly contemplative self- related (State 2) (MAAS: *r* = 0.549, *p* = 0.015; MWQ: *r* = 0.438, *p* = 0.056; RRQ_rumination: *r* = 0.426, *p* = 0.056). Given that MAAS measures the degree of attention and awareness in daily life, MWQ assesses MW tendencies, and RRQ_rumination evaluates rumination propensity, these findings suggest that the participants who have difficulty maintaining attention on ongoing activities in daily life show reduced awareness of physical and emotional changes, and are prone to mind wandering or becoming fixated on specific negative thought patterns, are more likely to experience attentional lapses and become immersed in self-related thoughts during experimental tasks.

### 3.2. Relationships Among Thought State Transitions, Autonomic Nervous Activity, and Interoceptive Accuracy

Before conducting the main analyses, we examined whether there were significant differences in autonomic nervous activity (RR intervals, RMSSD, and respiratory rate) across the identified thought states (State1,2,3). If any specific thought state exhibited a strong association with autonomic nervous activity, it would warrant a focused analysis of that state. However, linear mixed model estimation revealed no significant main effects of thought state for any of these measures (Appendix Fig. A1-3). These results suggest no significant differences in autonomic nervous activity associated with individual thought states. Additionally, to examine the validity of using the HCT to measure cardiac interoceptive accuracy, we calculated the correlation between the HCT and the TET scores. There was little correlation between the HCT and the TET scores (*r_32_* = 0.175, *p* = 0.337). Based on these results, the HCT scores are not significantly influenced by time estimation ability. Given this, in the following analyses, the average score of the six conditions of the HCT was used as an index of cardiac interoceptive accuracy. No significant correlations were observed between HCT scores and the occurrence probabilities of each thought state. From these results, we found no evidence that distinct patterns of autonomic nervous activity accompanied different thought states, nor that interoceptive characteristics explained the probability of experiencing specific thought states. Based on these results, we decided to investigate the relationship between sequential thought state transition patterns, autonomic nervous activity, and interoceptive accuracy rather than focusing on individual thought states in isolation.

Next, we examined the relationships among thought state transition patterns between consecutive trials during the task, autonomic nervous activity, and the participants’ interoceptive accuracy. We estimated multinomial linear regression models using thought state at trial t, RR intervals or RMSSD or respiratory rate, and individual HCT score as independent variables, and thought state at trial t+1 as the dependent variable. We focused on the interaction between thought state at trial t, autonomic indices, and interoceptive accuracy. The WAIC values were 555.9 for the RR interval model, 579.1 for the RMSSD model, and 582.6 for the respiratory rate model. After comparing the WAIC of these three models, we adopted the RR interval model, which showed the lowest WAIC. Information on each parameter in the other two models is provided in the Appendix (Tables A.1, A.2). Analysis of the RR interval model revealed that, compared to State1 (on-task)→2 (highly contemplative self-related) transitions, State2→2 transitions were more likely to occur, and compared to State1→3 (thought absence) transitions, State2→3 transitions and State3→3 continuation were more probable (State2→2: *b* = 2.474, 95% CI [1.166;3.813]; State2→3: *b* = 2.991, 95% CI [0.823;5.329]; State3→3: *b* =3.533, 95% CI [1.337;5.876]) (Table 1, Fig. 5A). These patterns are consistent with the thought state transition patterns observed in Section 3.1. Additionally, we found that the interaction between State2 (highly contemplative self-related thought), accompanying RR intervals, and individual interoceptive accuracy predicted the continuous of State2 in the subsequent trial (*b* = - 0.105, 95% CI [-0.24; −0.005]). Fig. 5B shows the relationship between thought transition pattern, RR, and HCT scores. The results indicate that when State2 was accompanied by decreased RR intervals (i.e., accelerated heart rate), the participants with higher cardiac interoceptive accuracy were more likely to continue State2 in the subsequent trial. As the 95% credible interval was largely outside the ROPE, this interaction was found to have a practical effect (Fig. 5A).

**Fig. 5.**
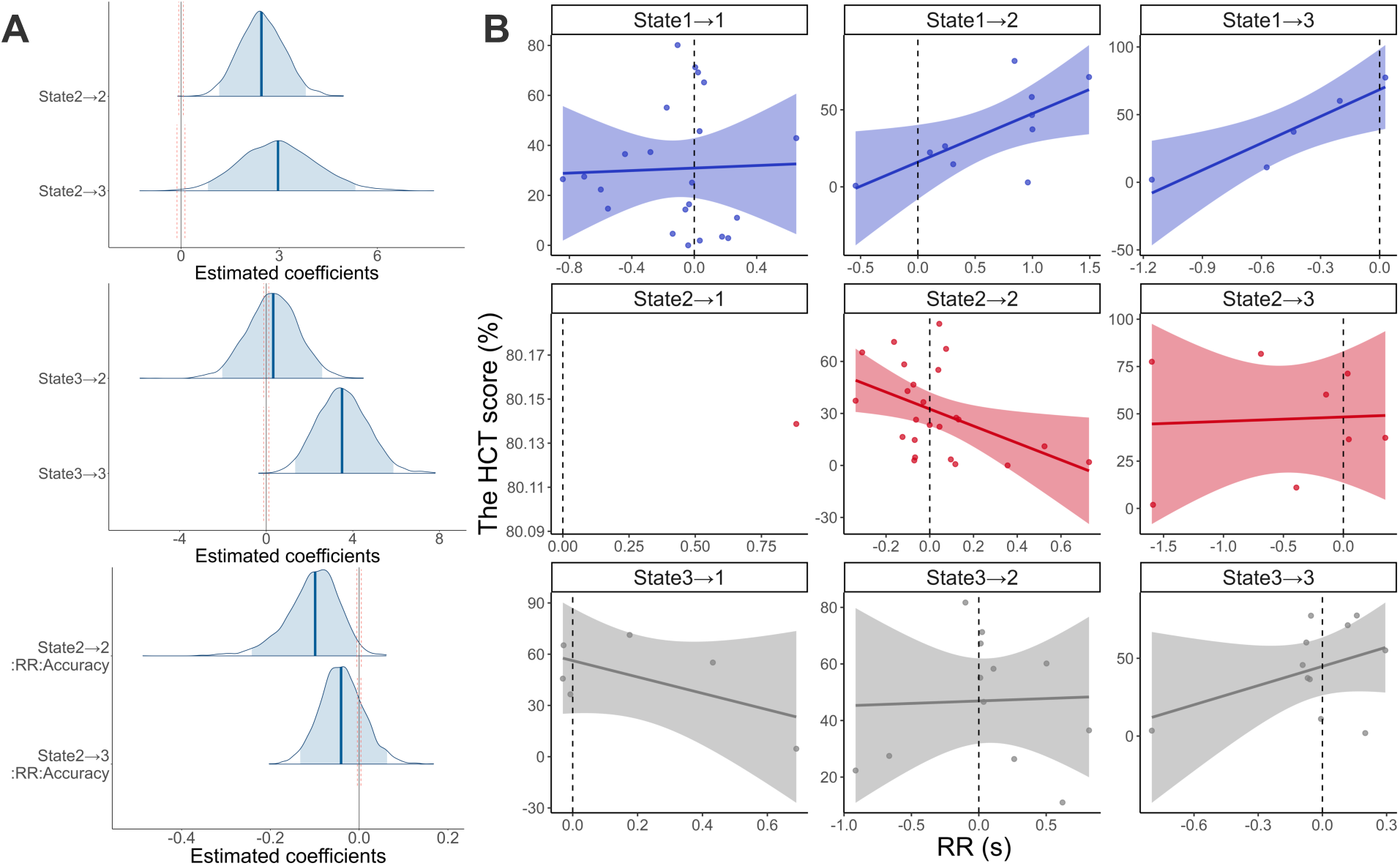
The Results of the Multinomial Linear Regression Model. (A) Distribution of the estimated parameters. The horizontal axis represents the estimated coefficients for each factor. Dark blue lines indicate the mean of the estimated coefficients, and the light blue ranges indicate 95 % credible intervals (CI). The orange wavy lines represent the region of practical equivalence (ROPE), the range within which a parameter has no practical effect. The 95 % CI of the estimated interactions among thought state, RR interval, and the score of the HCT largely exceeded the ROPE, revealing that these interactions had practical effects. (B) When State2 was accompanied by decreased RR intervals (i.e., accelerated heart rate), the participants with higher cardiac interoceptive accuracy were more likely to continue State2 in the subsequent trial.

**Table 1.** Summary of the estimated parameters in transition pattern of thought states.

| Parameters | Estimate | Est.Error | l-95% CI | u-95% CI | $\hat{R}$ | Bulk ESS | Tail ESS |
| --- | --- | --- | --- | --- | --- | --- | --- |
| State2_Intercept | -0.559 | 0.772 | -2.049 | 1.032 | 1 | 992.439 | 1794.072 |
| State3_Intercept | -4.765 | 1.205 | -7.302 | -2.580 | 1 | 1076.529 | 1715.459 |
| State2→2 | 2.474 | 0.680 | 1.166 | 3.813 | 1 | 1854.951 | 2680.249 |
| State3→2 | 0.299 | 1.183 | -2.034 | 2.585 | 1 | 1448.259 | 1535.154 |
| State2_RR | -0.276 | 0.739 | -1.759 | 1.198 | 1 | 1615.466 | 2051.886 |
| State2_Accuracy | -0.037 | 0.022 | -0.082 | 0.006 | 1 | 988.992 | 1563.892 |
| State2→2:RR | 1.479 | 1.215 | -0.900 | 4.002 | 1 | 1627.122 | 2556.975 |
| State3→2:RR | 1.571 | 2.244 | -2.807 | 6.124 | 1 | 1286.693 | 1246.685 |
| State2→2:Accuracy | 0.042 | 0.024 | -0.002 | 0.090 | 1 | 1175.336 | 1865.472 |
| State3→2:Accuracy | 0.046 | 0.028 | -0.005 | 0.100 | 1 | 1152.276 | 1627.089 |
| State2_RR:Accuracy | 0.062 | 0.049 | -0.015 | 0.182 | 1 | 878.795 | 910.084 |
| State2→2:RR:Accuracy | -0.105 | 0.060 | -0.242 | -0.005 | 1 | 841.533 | 935.725 |
| State3→2:RR:Accuracy | -0.110 | 0.104 | -0.338 | 0.068 | 1 | 904.632 | 1187.442 |
| State2→3 | 2.991 | 1.160 | 0.823 | 5.329 | 1 | 1287.345 | 1864.709 |
| State3→3 | 3.533 | 1.173 | 1.337 | 5.876 | 1 | 1205.031 | 1821.385 |
| State3_RR | -1.977 | 1.240 | -4.488 | 0.443 | 1 | 948.103 | 1780.164 |
| State3_Accuracy | 0.055 | 0.023 | 0.013 | 0.103 | 1 | 1000.396 | 1789.646 |
| State2→3:RR | 1.887 | 1.770 | -1.481 | 5.335 | 1 | 1101.555 | 1757.501 |
| State3→3:RR | -1.014 | 2.335 | -5.846 | 3.375 | 1 | 1237.472 | 1840.913 |
| State2→3:Accuracy | 0.000 | 0.026 | -0.052 | 0.050 | 1 | 1326.684 | 1812.713 |
| State3→3:Accuracy | -0.048 | 0.023 | -0.095 | -0.006 | 1 | 1303.653 | 1790.679 |
| State3_RR:Accuracy | 0.027 | 0.030 | -0.035 | 0.085 | 1 | 981.855 | 1447.806 |
| State2→3:RR:Accuracy | -0.039 | 0.049 | -0.133 | 0.063 | 1 | 1120.930 | 1605.689 |
| State3→3:RR:Accuracy | 0.024 | 0.060 | -0.096 | 0.144 | 1 | 1188.097 | 1534.470 |

Notes:
$\hat{R} = 1$ for all parameters, the model has converged.
Estimate: the estimated coefficient of each parameter.
Est. Error: the standard error in the estimated coefficient of each parameter.
l-95% CI: the lower limit of the 95% credible interval (CI).
u-95% CI: the upper limit of the 95% CI.
$\hat{R}$ : indicators for determining Markov Chain Monte Carlo (MCMC) chain convergence.
Usually, the model is considered to have converged if $\hat{R} \leq 1.1$ .
Bulk\_ESS: the adequate sample size calculated by normal rank.
Tail\_ESS: the adequate sample size for the 5% and 95% quantile points.

### 3.3. The Impact of Individual Characteristics on the Relationship Between Thought State Transitions, Autonomic Nervous Activity, and Interoceptive Accuracy

Furthermore, we comprehensively examined the relationships among individual differences in thought-related characteristics and experimentally measured thought transition patterns, autonomic nervous activity, and interoceptive accuracy. First, we computed Pearson’s correlation coefficients between the posterior predictive probability of each thought transition pattern and questionnaire scores (MWQ, DDFS, MAAS, STAI, BDI, RRQ_rumination, RRQ_reflection, BPQ). The analysis revealed significant negative correlations between BDI score and the predicted transition probability of State1 (on-task)→1 (*r* = −0.508, *p* = 0.007) and State1→2 (highly contemplative self-related; *r* = 0.445, *p* = 0.020). RRQ_rumination score correlated with the predicted transition probability of State1→1 (*r* = −0.499, *p* = 0.008), State1→2 (*r* = 0.493, *p* = 0.009) and State2→1 (*r* = −0.493, *p* = 0.012). MAAS score correlated with the predicted transition probability of State1→2 (*r* = 0.568, *p* = 0.002), State2→1 (*r* = −0.584, *p* = 0.002), State2→2 (*r* = 0.477, *p* = 0.016), and State3→2 (*r* = 0.632, *p* = 0.004). MWQ score correlated with the predicted transition probability of State1→2(*r* = 0.392, *p* = 0.043). These findings suggest that individuals with higher depressive tendencies and rumination are less likely to remain on-task (State1) and more likely to transition into highly contemplative self-related (State2). Additionally, the positive correlation with MAAS and MWQ suggests that individuals with lower present-moment awareness and higher MW tendencies are more prone to transition to State2.

For combinations of thought transition patterns and individual characteristics that showed significant correlations, we conducted structural equation modeling (SEM) to evaluate both direct and indirect pathways between individual characteristics and thought state transitions. In cases where a single indicator demonstrated significant correlations with multiple transition patterns, we considered transitions from the same initial state (e.g., State 1→1 and State 1→2) to represent trade-off relationships. Consequently, our modeling focused exclusively on transitions to task-unrelated thoughts, specifically State 2 or 3. Specifically, we examined (1) whether autonomic nervous activity (RR intervals) mediates these relationships and (2) whether interoceptive accuracy moderates the effects of autonomic nervous activity on thought transitions. The SEM results revealed several key findings in State1 (on-task)→2 (highly contemplative self-related) transitions (Table 2). First, BDI was positively associated with RR intervals (*β* = 0.045, *SE* = 0.018, *p* = 0.014) and interoceptive accuracy (*β* = 1.852, *SE* = 0.910, *p* = 0.042). These results suggest that higher depressive tendencies correspond to longer RR intervals and greater interoceptive accuracy. Regarding effects on State1→2 transition probability, RR intervals did not have a significant direct effect (*β* = −0.307, *SE* = 0.196, *p* = 0.118), whereas interoceptive accuracy showed a significant negative effect (*β* = −0.003, *SE* = 0.001, *p* = 0.003). Additionally, the interaction term (RR × Accuracy) had a significant effect on State1→2 transitions (*β* = 0.016, *SE* = 0.004, *p* < 0.001), indicating that the effect of RR intervals on on-task to self-related transitions was moderated by interoceptive accuracy. Regarding indirect effects, the mediation effect of RR intervals on the relationship between BDI and State1→2 was insignificant (Indirect effect = −0.014, *SE* = 0.012, *p* = 0.237). Similarly, the indirect effect through interoceptive accuracy was also non-significant (Indirect effect = −0.112, *SE* = 0.082, *p* = 0.172).

**Table 2.** Summary of the estimated parameters in the structural equation model of State1 (on-task)→2 (highly contemplative self-related) transition and BDI score

| Path | Estimate | Std.Error | Z.Value | P.Value | Lower | Upper |  |
| --- | --- | --- | --- | --- | --- | --- | --- |
| BDI⇒RR | 0.045 | 0.018 | 2.449 | 0.014 | 0.007 | 0.081 | ** |
| BDI⇒Interoceptive Accuracy | 1.852 | 0.910 | 2.034 | 0.042 | -0.166 | 3.393 | ** |
| BDI⇒State1→2 | 0.007 | 0.006 | 1.259 | 0.208 | -0.001 | 0.021 |  |
| RR⇒State1→2 | -0.307 | 0.196 | -1.564 | 0.118 | -0.713 | 0.066 |  |
| Interoceptive Accuracy⇒State1→2 | -0.003 | 0.001 | -3.015 | 0.003 | -0.005 | -0.001 | *** |
| RR: Interoceptive Accuracy⇒State1_2 | 0.016 | 0.004 | 4.025 | 0.000 | 0.008 | 0.025 | *** |
| Indirect effect RR | -0.014 | 0.012 | -1.183 | 0.237 | -0.044 | 0.003 |  |
| Indirect effect Interaction | 0.001 | 0.000 | 1.933 | 0.053 | 0.000 | 0.002 | * |
| Total indirect effect | -0.019 | 0.012 | -1.631 | 0.103 | -0.048 | 0.000 |  |
| Total effect | -0.012 | 0.012 | -0.992 | 0.321 | -0.040 | 0.010 |  |

\* $p < 0.1$ \*\* $p < 0.05$ \*\*\* $p < 0.01$
Notes:
Estimate: estimated path coefficient or effect within the model.
Std. Error: the standard error in the estimated value.
Lower: the lower limit of the 95% confidence interval.

However, the interaction-mediated indirect effect showed a trend-level effect but did not reach statistical significance (Indirect effect = 0.001, *SE* = 0.000, *p* = 0.053). The total indirect effect was estimated at −0.019 (*SE* = 0.012, *p* = 0.103). The total effect of BDI on State1→2 transition probability, including both direct and indirect pathways, was also non-significant (Total effect = −0.012, *SE* = 0.012, *p* = 0.321). These findings suggest that while depressive tendencies influence autonomic nervous activity (RR intervals) and interoceptive accuracy, the probability of transitioning from on-task to self-related is primarily determined by the interaction between RR intervals and interoceptive accuracy. This indicates that RR intervals alone are not a strong predictor of thought transitions, but their influence is contingent on interoceptive accuracy. The relationships between these variables are summarized in path diagrams in Fig. 6A. The relationships between State 1→2 transition probabilities and BDI scores, RR intervals, and interoceptive accuracy are summarized in Fig. 6B. For combinations other than BDI and State 1→2, no relationships were demonstrated where individual characteristic influenced thought state transition patterns through mediating effects on autonomic nervous activity or interoceptive accuracy. The results of these model analyses are summarized in the Appendix (Table A.3-A.7).

**Fig. 6.**
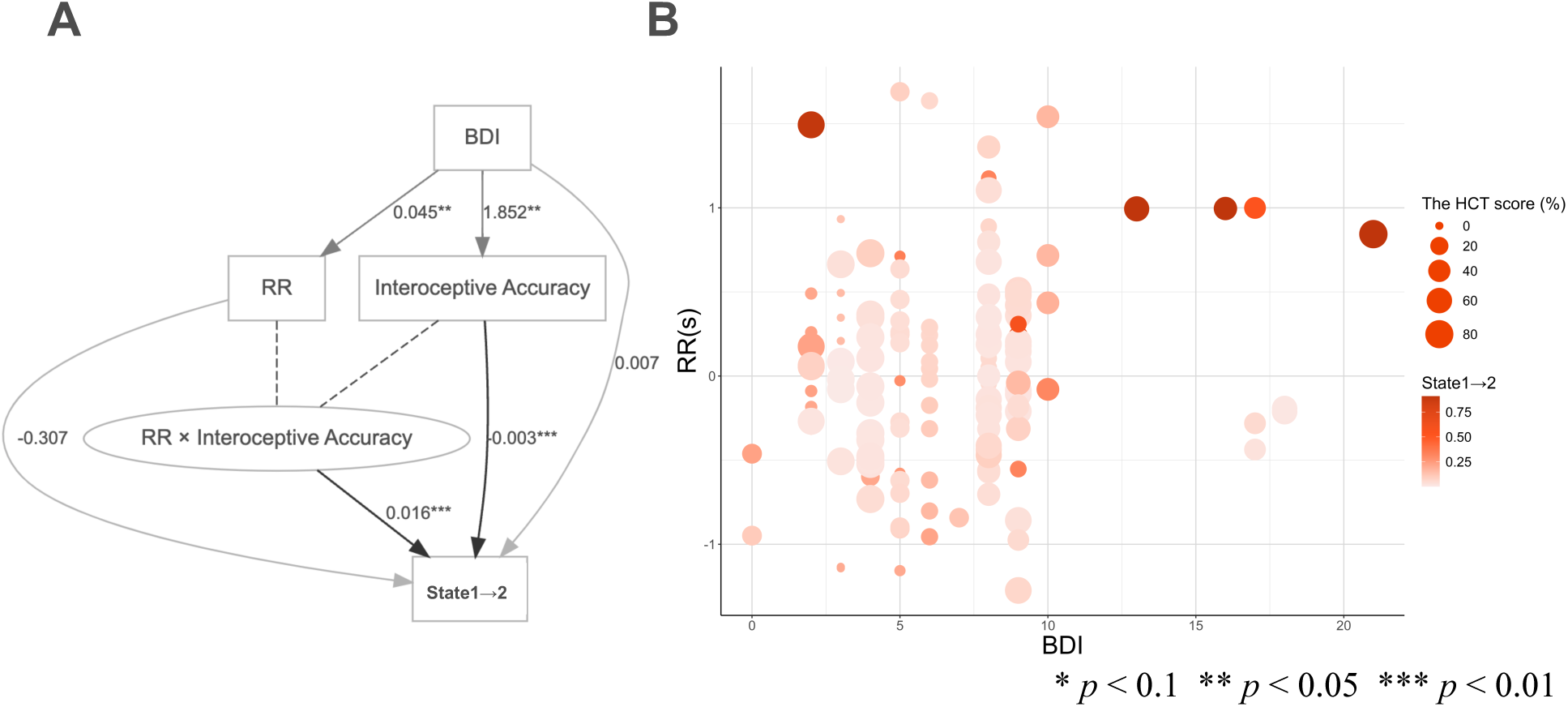
The Results of the Structural Equation Modeling. (A) Path diagram summarizing the mediation and moderation effects of RR intervals and interoceptive accuracy on the relationship between BDI scores and the probability of transition from State1 (on-task) to State2 (highly contemplative self-related). The figure illustrates the effects of BDI (Beck Depression Inventory) scores on RR intervals and interoceptive accuracy and their mediation and moderation effects on the probability of transitioning from State1 to State2. Dark gray lines represent pathways significant at the 1% level, medium gray lines represent pathways significant at the 5% level, and light gray lines represent non-significant pathways. The results suggest that BDI scores influence the probability of transitioning from State1 to State2 through the interaction between RR intervals and interoceptive accuracy. (B) Relationship between BDI scores, RR intervals, and interoceptive accuracy when a transition to State1→2 occurs. The x-axis represents BDI scores, the y-axis represents RR intervals, the size of the dots indicates HCT performance (i.e., interoceptive accuracy), and the color intensity of the dots reflects the probability of transitioning from State1 to State2. The results suggest that when both BDI scores and interoceptive accuracy are high, an increase in RR intervals (i.e., decreased heart rate) is associated with a higher probability of transitioning to State2.

## 4. Discussion

The present study has three objectives: (1) to estimate thought states by integrating multiple components of thought, (2) to examine the interaction among estimated thought state transitions, fluctuations in autonomic nervous activity, and cardiac interoception, and (3) to investigate how individual differences in depression, anxiety, daily tendencies in attention, and sensitivity to bodily sensations and emotions influence these relationships mediated by autonomic activity patterns and interoceptive accuracy. The participants periodically reported components of their thoughts while engaging in a simple attention task. Hidden Markov Models (HMM) were employed to estimate underlying thought states from time-series data of the participants’ thought components. We analyzed the relationships among thought state transition probabilities and patterns, task-related autonomic nervous activity, and interoceptive accuracy using multinomial logistic regression models. Furthermore, SEM was conducted to examine whether autonomic nervous activity and individual interoceptive accuracy mediated the influence of thought-related individual differences on thought transition patterns. Based on our previous findings (Sakuragi et al., 2023, 2024), we hypothesized that (1) sympathetic dominance would promote the persistence of self-related thoughts, while parasympathetic dominance would facilitate thought transitions, and (2) the influence of individual differences in thought-related characteristics on thought transition patterns would be mediated by real-time autonomic activity and further modulated by interoceptive accuracy.

The HMM analysis revealed that the participants’ thoughts during task performance could be classified into three states (Fig. 4): on-task (State 1), highly contemplative self-related (State 2), and thought absence (State 3). State 1 had low arousal and contemplation levels with neutral emotional valence. In contrast, State 2 exhibited high arousal and contemplation levels, often accompanied by strong negative or positive emotions. These results suggest that self-related thoughts occurring during a simple attention task tend to be characterized by higher levels of arousal and contemplation and are more likely to be accompanied by richer emotional valence than task-focused states.

However, the thought states extracted at each time point through HMM analysis did not simply correspond to different autonomic nervous activities. Instead, fluctuations in autonomic nervous activity during the task interacted with the participants’ interoceptive accuracy and their current thought state, subsequently influencing their thought state at the next time point. More specifically, the participants with high interoceptive accuracy were more likely to maintain State 2 (highly contemplative self-related) in subsequent trials when self-related thoughts were accompanied by increased heart rate (shorter RR intervals). This suggests that when the activation of sympathetic nervous activity accompanies self-related thoughts, accurately detecting this activation may facilitate deeper engagement in self-related thoughts. This immersion in self-related states may be underpinned by the activation of first-person conscious states associated with updating interoceptive information triggered by physiological arousal accompanying self-related trial states. The results of the present study align with our previous research and support our hypothesis regarding the role of sympathetic activation in the maintenance of self-related thoughts.

Notably, the probability of reporting State 2 showed a moderate positive correlation with individual rumination tendencies. Indeed, State 2 was characterized by high arousal and deep contemplation, often accompanied by strongly negative emotions, aligning with the characteristics of rumination. Furthermore, spontaneous thoughts involving rumination, worry, and anxiety have been associated with sympathetic nervous activation (i.e., decreased vagal activity; Ottaviani et al., 2009, 2016; Ottaviani, Shahabi, et al., 2015).

Considering these factors, it is possible that part of the sustained presence of State 2, which was shown to interact with autonomic nervous activity and interoceptive accuracy, reflects a ruminative state during the experimental task.

In contrast to the maintenance of State 2, the transition from State 1 (on-task) to State 2 (highly contemplative self-related) was influenced by different factors. The SEM analysis revealed that the probability of transitioning from State 1 to State 2 was significantly influenced by the interaction between interoceptive accuracy and autonomic nervous activity, but only among individuals with higher depressive tendencies (BDI scores). Specifically, when parasympathetic nervous activity was dominant (longer RR intervals), individuals with both high depressive tendencies and high interoceptive accuracy were more likely to transition from State 1 to State 2. These findings suggest that interoceptive accuracy plays distinct roles in thought dynamics: it facilitates the persistence of self-related thoughts when sympathetic activation is high, whereas, in the presence of depressive tendencies, it increases the likelihood of transitioning to self-related thoughts under parasympathetic dominance.

Building upon the discussion in the previous paragraph, assuming State 2 encompasses a ruminative-like state frequently observed in depressive conditions, the transition from State 1 to State 2 may reflect the initiation of rumination during the experimental task. The activation of parasympathetic activity at the onset of rumination may be explained by the involvement of the default mode network (DMN) in both self-referential thinking or depressive rumination and parasympathetic regulation (Beissner et al., 2013; Hamilton et al., 2015; Zhou et al., 2020). This suggests that during the initiation of mind wandering, DMN activation may trigger parasympathetic activity, which, when accurately detected through interoceptive processes, could potentially trigger rumination. Future research should investigate whether DMN activation and parasympathetic activity occur sequentially during transitions into rumination, followed by the activation of interoceptive neural processing. If this hypothesis proves correct, measuring interoceptive accuracy and monitoring autonomic nervous activity in individuals prone to rumination could contribute to developing systems that predict deep engagement into ruminative states before they fully manifest.

One limitation of the present study is that we did not thoroughly examine the detailed characteristics of thought components, such as temporal orientation and emotional valence, in relation to thought transition patterns, autonomic nervous activity, and interoceptive accuracy. While we would expect specific patterns to emerge, mainly about past negative thought states, if the phenomenon observed in this study reflects rumination, our data did not yield such results. The potential differences in temporal orientation and emotional valence between or within individuals may have been diminished due to our methodology of varying questions across thought categories, retrospectively supplementing data, and requesting continuous responses for all items. Based on these findings, future studies should consider employing Likert scales and devise ways to collect subjective data from participants at as many time points as possible. Another limitation is that the causal relationships among thought transitions, autonomic nervous activity, and interoceptive accuracy remain unclear. Future studies should employ time-series causal models or dynamic Bayesian networks to better capture these factors’ dynamic and causal relationships.

## 5. Conclusion

This study aimed to investigate the relationships among thought state transitions, autonomic nervous activity, and interoception, as well as the influence of individual difference factors on these interactions. The results revealed that (1) higher interoceptive accuracy facilitated the persistence of emotionally charged, highly contemplative self-referential thoughts under sympathetic dominance, and (2) individuals with higher depressive tendencies were more likely to transition into self-referential thoughts during parasympathetic dominance, with this effect being modulated by interoceptive accuracy. These findings suggest that spontaneous thoughts emerge from a complex interplay between thought-related individual characteristics, real-time autonomic nervous activity, and interoceptive accuracy. This study provides important insights into the relationship between mental health and physiological responses, particularly in understanding thought flexibility and fixation. Moreover, it may contribute to a better understanding of psychiatric disorders characterized by excessive rumination and reduced attentional flexibility. Future research should carefully examine whether similar autonomic and interoceptive contributions are observed not only in transient rumination among healthy individuals but also in the pathological and persistent rumination seen in major depressive and anxiety disorders.

## Ethics approval

The study was approved by the Keio University Research Ethics Committee (No. 240120000), Japan.

## Consent to participate

All participants gave informed consent to participate in the present study, and their rights were protected.

## Availability of data and materials

The datasets analyzed during this study are available from the corresponding author upon reasonable request.

## CRediT authorship contribution statement

**Mai Sakuragi:** Writing – original draft, Software, Methodology, Investigation, Formal analysis, Conceptualization, Funding acquisition. **Satoshi Umeda:** Writing – review & editing, Supervision.

## Acknowledgments

This work was supported by JSPS KAKENHI Grant Number JP24KJ1954.

