## Appendix for "How the Heart Shapes the Mind: The Role of Cardiac Interoception in the Interaction between Autonomic Nervous Activity and Self-related Thoughts"

Contents of the original questionnaire about participants’ thought patterns in daily life during the week before the experiment.

Participants responded to the following questions: (1) During the past week, how often did specific episodes or images spontaneously come to mind? (1. Never, 2. Sometimes (several times per day), 3. Frequently (about once every few hours), 4. Constantly (about once every few minutes)), (2) Which type of episodes or images were most prevalent in your mind during the past week? (1. Personal past experiences, 2. Personal future plans, 3. Knowledge or facts from conversations with others, TV, radio, social media, newspapers, or books, 4. Images or fantasies related to books, videos, music, or artwork that you experienced), (3) Regarding the episodes or images reported in (2), rate the emotional valence and arousal level of the episodes or images themselves (1. Very negative to 5. Very positive / 1. Very calm to 5. Very excited), (4) Regarding the episodes or images reported in (2), rate the emotional valence and arousal level of yourself while recalling these episodes or images (1. Very negative to 5. Very positive / 1. Very calm to 5. Very excited), and (5) To what extent were the episodes or images reported in (2) recalled during the experimental task? (1. Not at all, 2. Sometimes (2-5 times), 3. Frequently (6-9 times), 4. Constantly (10 times or more)). If participants selected "Never" for question (1), the subsequent questions (2-5) were not presented.

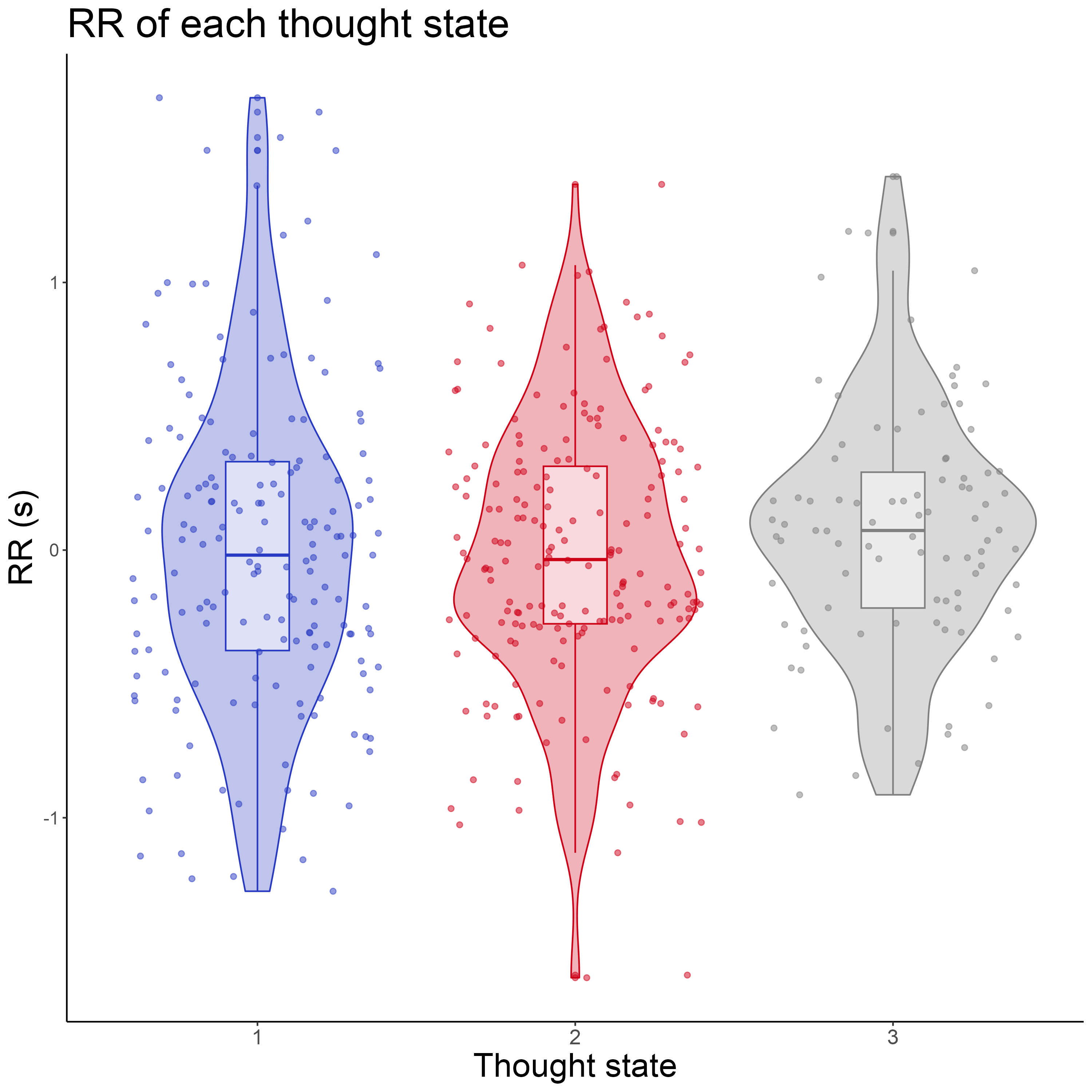

Fig. A.1. RR interval for each thought state

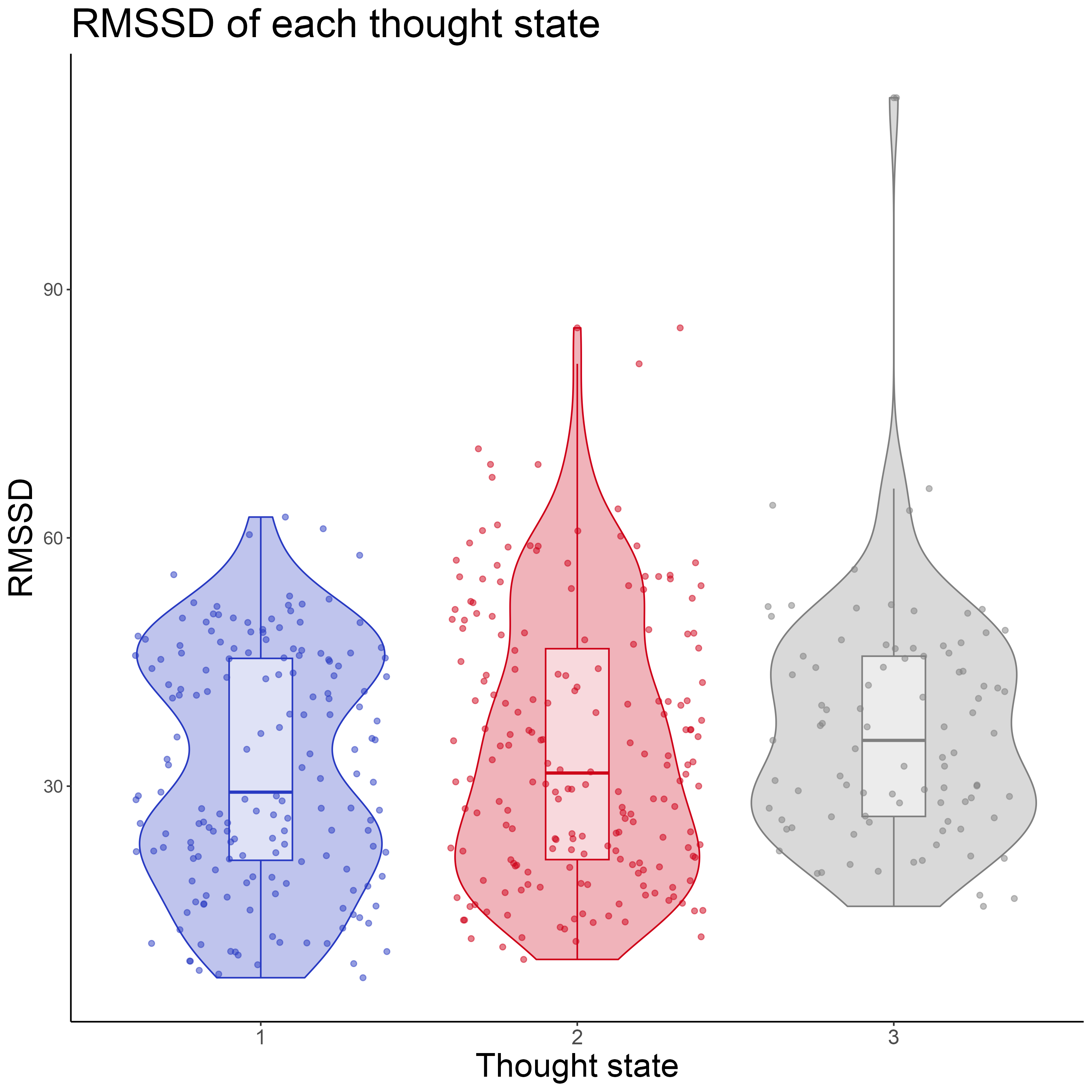

Fig. A.2. RMSSD interval for each thought state

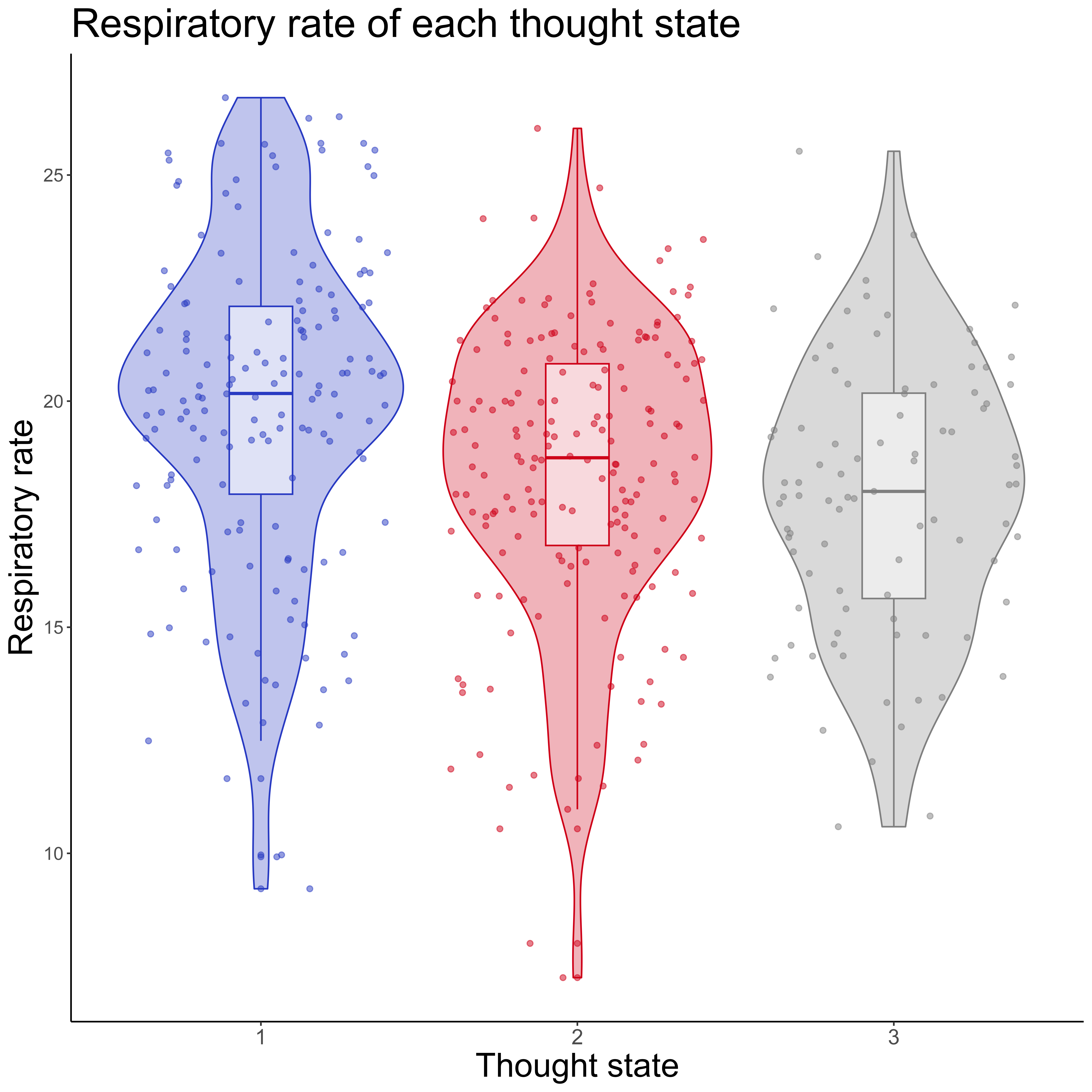

Fig. A.3. Respiratory rate interval for each thought state

Table A.1. Summary of the estimated parameters in transition pattern of thought states (RMSSD ver.).

| Parameters | Estimate | Est.Error | l-95% CI | u-95% CI | R̂ | Bulk_ESS | Tail_ESS |
| --- | --- | --- | --- | --- | --- | --- | --- |
| State2_Intercept | -2.768 | 1.550 | -6.035 | 0.025 | 1 | 1505.581 | 2494.231 |
| State3_Intercept | -7.387 | 2.561 | -12.762 | -2.576 | 1 | 1347.255 | 1960.597 |
| State2→2 | 1.441 | 1.595 | -1.635 | 4.507 | 1 | 1999.103 | 1921.629 |
| State3→2 | 2.759 | 3.493 | -4.070 | 9.824 | 1 | 1535.216 | 1516.316 |
| State2_RMSSD | 0.071 | 0.054 | -0.025 | 0.182 | 1 | 1085.749 | 1631.764 |
| State2_Accuracy | 0.017 | 0.062 | -0.102 | 0.143 | 1 | 1620.407 | 1684.456 |
| State2→2:RMSSD | 0.045 | 0.060 | -0.067 | 0.167 | 1 | 1676.549 | 2081.880 |
| State3→2:RMSSD | -0.056 | 0.094 | -0.242 | 0.124 | 1 | 1433.458 | 1782.774 |
| State2→2:Accuracy | 0.056 | 0.070 | -0.082 | 0.192 | 1 | 1834.754 | 2318.360 |
| State3→2:Accuracy | -0.017 | 0.094 | -0.208 | 0.168 | 1 | 1413.472 | 1660.202 |
| State2_RMSSD:Accuracy | -0.002 | 0.002 | -0.007 | 0.002 | 1 | 1257.783 | 1177.205 |
| State2→2:RMSSD:Accuracy | 0.000 | 0.003 | -0.005 | 0.005 | 1 | 1454.592 | 1362.711 |
| State3→2:RMSSD:Accuracy | 0.002 | 0.003 | -0.004 | 0.009 | 1 | 1348.885 | 1367.031 |
| State2→3 | 6.047 | 2.926 | 0.650 | 11.865 | 1 | 1077.076 | 1684.443 |
| State3→3 | 7.007 | 4.076 | -0.731 | 15.377 | 1 | 1264.818 | 1703.482 |
| State3_RMSSD | 0.094 | 0.073 | -0.058 | 0.236 | 1 | 1316.931 | 1900.871 |
| State3_Accuracy | 0.187 | 0.077 | 0.049 | 0.356 | 1 | 1155.647 | 1324.140 |
| State2→3:RMSSD | -0.096 | 0.089 | -0.271 | 0.071 | 1 | 1076.322 | 1923.444 |
| State3→3:RMSSD | -0.132 | 0.106 | -0.347 | 0.066 | 1 | 1140.375 | 1681.212 |
| State2→3:Accuracy | -0.152 | 0.092 | -0.351 | 0.015 | 1 | 1062.688 | 1306.170 |
| State3→3:Accuracy | -0.173 | 0.104 | -0.392 | 0.020 | 1 | 1080.677 | 1344.982 |
| State3_RMSSD:Accuracy | -0.004 | 0.002 | -0.009 | 0.000 | 1 | 1119.356 | 1394.650 |
| State2→3:RMSSD:Accuracy | 0.005 | 0.003 | -0.001 | 0.011 | 1 | 1022.992 | 1516.174 |
| State3→3:RMSSD:Accuracy | 0.004 | 0.003 | -0.001 | 0.011 | 1 | 1024.287 | 1492.031 |

Table A.2. Summary of the estimated parameters in transition pattern of thought states (Respiratory rate ver.).

| Parameters | Estimate | Est.Error | l-95% CI | u-95% CI | R̂ | Bulk_ESS | Tail_ESS |
| --- | --- | --- | --- | --- | --- | --- | --- |
| State2_Intercept | 4.712 | 3.700 | -2.295 | 12.479 | 1 | 1768.820 | 2341.169 |
| State3_Intercept | -0.469 | 4.774 | -10.529 | 8.224 | 1 | 1516.139 | 1973.982 |
| State2→2 | -2.530 | 6.013 | -14.046 | 9.203 | 1 | 1391.334 | 2268.617 |
| State3→2 | 2.209 | 8.492 | -14.054 | 19.540 | 1 | 1760.406 | 2713.380 |
| State2_Resp | -0.281 | 0.185 | -0.668 | 0.073 | 1 | 1814.620 | 2422.299 |
| State2_Accuracy | -0.121 | 0.142 | -0.419 | 0.147 | 1 | 1607.964 | 2061.870 |
| State2→2:Resp | 0.278 | 0.302 | -0.313 | 0.861 | 1 | 1362.787 | 2275.426 |
| State3→2:Resp | -0.072 | 0.453 | -1.000 | 0.796 | 1 | 1766.689 | 2516.037 |
| State2→2:Accuracy | 0.440 | 0.242 | -0.013 | 0.940 | 1 | 1297.419 | 1660.912 |
| State3→2:Accuracy | 0.114 | 0.213 | -0.288 | 0.547 | 1 | 1490.270 | 1930.404 |
| State2_Resp:Accuracy | 0.004 | 0.008 | -0.011 | 0.019 | 1 | 1627.813 | 2235.735 |
| State2→2:Resp:Accuracy | -0.020 | 0.012 | -0.045 | 0.004 | 1 | 1264.796 | 1591.284 |
| State3→2:Resp:Accuracy | -0.004 | 0.012 | -0.027 | 0.019 | 1 | 1511.033 | 2210.975 |
| State2→3 | -0.270 | 7.460 | -14.255 | 14.882 | 1 | 1315.028 | 2233.673 |
| State3→3 | 4.003 | 8.484 | -11.999 | 21.786 | 1 | 1506.410 | 1881.406 |
| State3_Resp | -0.147 | 0.237 | -0.592 | 0.351 | 1 | 1519.894 | 1932.575 |
| State3_Accuracy | -0.003 | 0.141 | -0.274 | 0.293 | 1 | 1448.942 | 1703.650 |
| State2→3:Resp | 0.133 | 0.382 | -0.633 | 0.865 | 1 | 1312.964 | 2204.560 |
| State3→3:Resp | -0.094 | 0.449 | -1.039 | 0.757 | 1 | 1515.995 | 1919.501 |
| State2→3:Accuracy | 0.403 | 0.249 | -0.065 | 0.916 | 1 | 1184.135 | 1955.127 |
| State3→3:Accuracy | 0.006 | 0.193 | -0.394 | 0.380 | 1 | 1263.495 | 1513.349 |
| State3_Resp:Accuracy | 0.001 | 0.008 | -0.015 | 0.015 | 1 | 1474.451 | 1618.716 |
| State2→3:Resp:Accuracy | -0.019 | 0.013 | -0.046 | 0.005 | 1 | 1168.755 | 1969.900 |
| State3→3:Resp:Accuracy | 0.000 | 0.010 | -0.021 | 0.021 | 1 | 1277.484 | 1605.418 |

Table A.3. Summary of the estimated parameters in the structural equation model of State1→2 transition and RRQ_rumination score

| Path | Estimate | Std.Error | Z.Value | P.Value | Lower | Upper |  |
| --- | --- | --- | --- | --- | --- | --- | --- |
| RRQ ⇨RR | 0.016 | 0.011 | 1.382 | 0.167 | -0.007 | 0.037 |  |
| RRQ ⇨Interoceptive Accuracy | 0.536 | 0.755 | 0.709 | 0.478 | -1.167 | 1.763 |  |
| RRQ ⇨State1→2 | 0.008 | 0.005 | 1.668 | 0.095 | -0.001 | 0.017 | * |
| RR⇨State1→2 | -0.244 | 0.202 | -1.210 | 0.226 | -0.657 | 0.121 |  |
| Interoceptive Accuracy⇨State1→2 | -0.003 | 0.001 | -2.750 | 0.006 | -0.005 | -0.001 | *** |
| RR:Interoceptive Accuracy⇨State1→2 | 0.015 | 0.004 | 3.565 | 0.000 | 0.008 | 0.024 | *** |
| Indirect effect RR | -0.004 | 0.005 | -0.735 | 0.462 | -0.016 | 0.004 |  |
| Indirect effect Interaction | 0.000 | 0.000 | 1.168 | 0.243 | 0.000 | 0.001 |  |
| Total indirect effect | -0.005 | 0.006 | -0.864 | 0.388 | -0.020 | 0.005 |  |
| Total effect | 0.003 | 0.008 | 0.366 | 0.714 | -0.014 | 0.017 |  |

* *p* < 0.1 ** *p* < 0.05 *** *p* < 0.01

Table A.4. Summary of the estimated parameters in the structural equation model of State1→2 transition and MAAS score

| Path | Estimate | Std.Error | Z.Value | P.Value | Lower | Upper |  |
| --- | --- | --- | --- | --- | --- | --- | --- |
| MAAS⇨RR | 0.006 | 0.007 | 0.863 | 0.388 | -0.008 | 0.018 |  |
| MAAS⇨Interoceptive Accuracy | -0.415 | 0.468 | -0.888 | 0.375 | -1.405 | 0.457 |  |
| MAAS⇨State1→2 | 0.007 | 0.003 | 2.115 | 0.034 | 0.001 | 0.013 | ** |
| RR⇨State1→2 | -0.244 | 0.163 | -1.498 | 0.134 | -0.610 | 0.017 |  |
| Interoceptive Accuracy⇨State1→2 | -0.002 | 0.001 | -1.478 | 0.139 | -0.004 | 0.001 |  |
| RR:Interoceptive Accuracy⇨State1→2 | 0.015 | 0.004 | 3.980 | 0.000 | 0.007 | 0.022 | *** |
| Indirect effect RR | -0.001 | 0.002 | -0.654 | 0.513 | -0.006 | 0.003 |  |
| Indirect effect Interaction | 0.000 | 0.000 | 0.835 | 0.404 | 0.000 | 0.000 |  |
| Total indirect effect | -0.001 | 0.003 | -0.194 | 0.846 | -0.006 | 0.005 |  |
| Total effect | 0.006 | 0.004 | 1.745 | 0.081 | -0.002 | 0.012 |  |

* *p* < 0.1 ** *p* < 0.05 *** *p* < 0.01

Table A.5. Summary of the estimated parameters in the structural equation model of State2→2 transition and MAAS score

| Path | Estimate | Std.Error | Z.Value | P.Value | Lower | Upper |  |
| --- | --- | --- | --- | --- | --- | --- | --- |
| MAAS⇨RR | -0.009 | 0.006 | -1.587 | 0.112 | -0.021 | 0.002 |  |
| MAAS⇨Interoceptive Accuracy | -0.376 | 0.513 | -0.733 | 0.464 | -1.397 | 0.611 |  |
| MAAS⇨State2→2 | 0.015 | 0.003 | 4.264 | 0.000 | 0.008 | 0.022 | *** |
| RR⇨State2→2 | 1.111 | 0.647 | 1.716 | 0.086 | -0.148 | 2.419 | * |
| Interoceptive Accuracy⇨State2→2 | 0.000 | 0.002 | -0.040 | 0.968 | -0.004 | 0.004 |  |
| RR:Interoceptive ⇨State2→2 | -0.011 | 0.010 | -1.170 | 0.242 | -0.031 | 0.006 |  |
| Indirect effect RR | -0.010 | 0.010 | -1.021 | 0.307 | -0.036 | 0.003 |  |
| Indirect effect Interaction | 0.000 | 0.000 | 0.849 | 0.396 | 0.000 | 0.000 |  |
| Total indirect effect | -0.010 | 0.010 | -1.002 | 0.316 | -0.035 | 0.003 |  |
| Total effect | 0.005 | 0.008 | 0.554 | 0.580 | -0.017 | 0.016 |  |

* *p* < 0.1 ** *p* < 0.05 *** *p* < 0.01

Table A.6. Summary of the estimated parameters in the structural equation model of State3→2 transition and MAAS score

| Path | Estimate | Std.Error | Z.Value | P.Value | Lower | Upper |  |
| --- | --- | --- | --- | --- | --- | --- | --- |
| MAAS⇨RR | 0.005 | 0.005 | 0.943 | 0.346 | -0.007 | 0.014 |  |
| MAAS⇨Interoceptive Accuracy | -0.531 | 0.408 | -1.303 | 0.192 | -1.297 | 0.303 |  |
| MAAS⇨State3→2 | 0.019 | 0.003 | 5.840 | 0.000 | 0.012 | 0.025 | *** |
| RR⇨State3→2 | -0.167 | 0.378 | -0.442 | 0.659 | -0.866 | 0.488 |  |
| Interoceptive Accuracy⇨State3→2 | 0.005 | 0.002 | 2.667 | 0.008 | 0.001 | 0.009 | *** |
| RR:Interoceptive Accuracy⇨State3→2 | -0.012 | 0.012 | -0.975 | 0.329 | -0.036 | 0.015 |  |
| Indirect effect RR | -0.001 | 0.002 | -0.396 | 0.692 | -0.005 | 0.003 |  |
| Indirect effect Interaction | 0.000 | 0.000 | -0.623 | 0.533 | 0.000 | 0.000 |  |
| Total indirect effect | -0.004 | 0.003 | -1.182 | 0.237 | -0.011 | 0.001 |  |
| Total effect | 0.015 | 0.004 | 4.317 | 0.000 | 0.009 | 0.023 | *** |

* *p* < 0.1 ** *p* < 0.05 *** *p* < 0.01

Table A.7. Summary of the estimated parameters in the structural equation model of State1→2 transition and MWQ score

| Path | Estimate | Std.Error | Z.Value | P.Value | Lower | Upper |  |
| --- | --- | --- | --- | --- | --- | --- | --- |
| MWQ⇨RR | 0.001 | 0.067 | 0.022 | 0.982 | -0.150 | 0.121 |  |
| MWQ⇨Interoceptive Accuracy | -6.886 | 3.737 | -1.843 | 0.065 | -14.089 | 0.277 | * |
| MWQ⇨State1→2 | 0.072 | 0.026 | 2.733 | 0.006 | 0.015 | 0.117 | *** |
| RR⇨State1→2 | -0.318 | 0.186 | -1.704 | 0.088 | -0.676 | 0.065 | * |
| Interoceptive Accuracy⇨State1→2 | -0.002 | 0.001 | -1.513 | 0.130 | -0.004 | 0.001 |  |
| RR:Interoceptive Accuracy⇨State1→2 | 0.017 | 0.004 | 4.132 | 0.000 | 0.009 | 0.025 | *** |
| Indirect effect RR | 0.000 | 0.028 | -0.017 | 0.987 | -0.048 | 0.071 |  |
| Indirect effect Interaction | 0.000 | 0.001 | 0.020 | 0.984 | -0.003 | 0.002 |  |
| Total indirect effect | 0.011 | 0.032 | 0.344 | 0.731 | -0.050 | 0.095 |  |
| Total effect | 0.083 | 0.028 | 2.928 | 0.003 | 0.026 | 0.137 | *** |

* *p* < 0.1 ** *p* < 0.05 *** *p* < 0.01
